# Parsing multiomics landscape of activated synovial fibroblasts highlights drug targets linked to genetic risk of rheumatoid arthritis

**DOI:** 10.1101/861781

**Authors:** Haruka Tsuchiya, Mineto Ota, Shuji Sumitomo, Kazuyoshi Ishigaki, Akari Suzuki, Toyonori Sakata, Yumi Tsuchida, Hiroshi Inui, Jun Hirose, Yuta Kochi, Yuho Kadono, Katsuhiko Shirahige, Sakae Tanaka, Kazuhiko Yamamoto, Keishi Fujio

## Abstract

**Objectives:** Synovial fibroblasts (SFs) produce a variety of pathogenic molecules in the inflamed synovium of rheumatoid arthritis (RA). We aimed to gain insight into the pathogenic mechanisms of SFs through elucidating the genetic contribution to molecular regulatory networks under inflammatory condition.

**Methods:** SFs from RA and osteoarthritis (OA) patients (n=30 each) were stimulated with 8 different cytokines (IFN-α, IFN-γ, TNF-α, IL-1β, IL-6/sIL-6R, IL-17, TGF-β1, IL-18) or a combination of all 8 (8-mix). Peripheral blood mononuclear cells (PBMCs) from the same patients were fractioned into five major immune cell subsets (CD4^+^ T cells, CD8^+^ T cells, B cells, NK cells, monocytes). Integrative analyses including mRNA expression, histone modifications (H3K27ac, H3K4me1, H3K4me3), 3D genome architecture and genetic variations of SNPs were performed.

**Results:** SFs exposed to synergistically acting cytokines produced markedly higher levels of pathogenic molecules, including CD40 whose expression was significantly affected by a RA risk SNP (rs6074022). Upon chromatin remodeling in activated SFs, RA risk loci were enriched in clusters of enhancers (super-enhancers; SEs) induced by synergistic proinflammatory cytokines. A RA risk SNP (rs28411362), located in a SE under synergistically acting cytokines, formed three-dimensional contact with the promoter of *MTF1* gene, whose binding motif showed significant enrichment in stimulation specific-SEs. Consistently, inhibition of MTF1 suppressed cytokine and chemokine production from SFs and ameliorated mice model of arthritis.

**Conclusions:** Our findings established the dynamic landscape of activated SFs, and yielded potential therapeutic targets associated with genetic risk of RA.

**Key messages:** *What is already known about this subject?:*

- In rheumatoid arthritis (RA), a variety of dysregulated molecules from immune cells and mesenchymal cells drive disease progression. Synovial fibroblasts (SFs), the most abundant resident mesenchymal cells in the inflamed synovium, produce a variety of pathogenic molecules including IL-6.
- Genome-wide association studies (GWAS) have identified more than 100 RA susceptibility loci. To gain insight into the pathogenic mechanisms of SFs, understanding the genetic contribution to molecular regulatory networks under inflammatory condition is crucial.

*What does this study add?:*

- Integrated analyses of activated SFs demonstrated that SFs exposed to synergistically acting cytokines produced markedly higher levels of pathogenic molecules. Some of which were significantly affected by RA risk loci in a stimulation-specific manner.
- Chromatin remodeling induced by synergistic proinflammatory cytokines were associated with RA heritability. Some transcription factors (MTF1, RUNX1) could be crucial for this structural rearrangement and the formation of inflammatory arthritis.

*How might this impact on clinical practice or future developments?:*

- Our findings established the dynamic landscape of activated SFs, and yielded potential therapeutic targets associated with genetic risk of RA.

## Introduction

Rheumatoid arthritis (RA) causes persistent synovitis leading to disabling joint destruction. Current treatment strategies that target cytokines (e.g., TNF-α, IL-6), cell surface proteins (e.g., CD20, CD80/86) or signaling molecules (e.g., Janus kinase) have brought a paradigm shift in RA treatment. However, achieving sustained remission is still challenging even with such agents.[1] Although the concepts of targeting multiple molecules have been proposed, combination or bispecific antibodies (anti-TNF-α and anti-IL-1β or anti-IL-17) failed to improve therapeutic efficacy.[2] These findings imply that some unknown factors play critical roles in the progression of synovitis.

In the pathogenesis of RA, the activities of a variety of dysregulated molecules in immune cells and mesenchymal cells are orchestrated by genetic and environmental factors.[3] To date, more than 100 RA susceptibility loci have been identified in genome-wide association studies (GWAS).[4] Recent genetic studies of autoimmune diseases have reported that the majority (> 90%) of these risk variants are located in non-coding regions and regulate gene expression in a cell type-specific manner,[5] partly in an environment-specific fashion.[6] An integrated understanding of the risk variants’ contribution to gene regulatory networks is crucial to gain insight into the pathogenic mechanisms of RA.

Synovial fibroblasts (SFs), the most abundant resident mesenchymal cells in the synovium, are major local effectors in the initiation and perpetuation of destructive joint inflammation through their production of a variety of pathogenic molecules including IL-6.[3] Previous multiomics data of unstimulated SFs have proposed activated pathways in RASFs.[7] However a comprehensive picture of SFs’ contribution to RA pathogenesis has largely remained elusive, perhaps due to their complex features that mutate in response to the proinflammatory milieu.[8] To date, a number of single cytokines that induce the inflammatory behavior of SFs have been reported (e.g., IFN-γ and IL-17 from T cells and TNF-α, IL-1β, IFN-α, IL-18 and TGF-β1 from monocytes).[9] To make matters more complicated, in *in vivo*, SFs are expected to be exposed to a more complex environment. Data show that some cytokine combinations (e.g., TNF-α and IL-17) synergistically enhance the expression of cytokines and chemokines.[10] Those findings emphasize the need to analyze the mechanisms underlying the accelerated inflammatory behavior of SFs in the presence of synergistic factors.

Here, we used integrative methods to analyze genomic, transcriptomic and epigenomic features of RASFs in the presence of various proinflammatory cytokines in RA joints.[11] We demonstrate that SFs exposed to synergistic proinflammatory cytokines show distinct transcriptomic features characterized by elevated expression of pathogenic molecules accompanied by chromatin remodeling associated with genetic risk of RA.

## Materials and methods

See online supplementary materials and methods.

## Results

### Cytokine mixture induce a distinctive transcriptome signature in SFs

We stimulated SFs from RA and osteoarthritis (OA) patients (n=30 each) with 8 different cytokines (IFN-α, IFN-γ, TNF-α, IL-1β, IL-6/sIL-6R, IL-17, TGF-β1, IL-18) or a combination of all 8 (8-mix). We also fractionated peripheral blood mononuclear cells (PBMCs) from the same patients into five major immune cell subsets (CD4^+^ T cells, CD8^+^ T cells, B cells, NK cells, monocytes) (figure 1).

**Figure 1.**
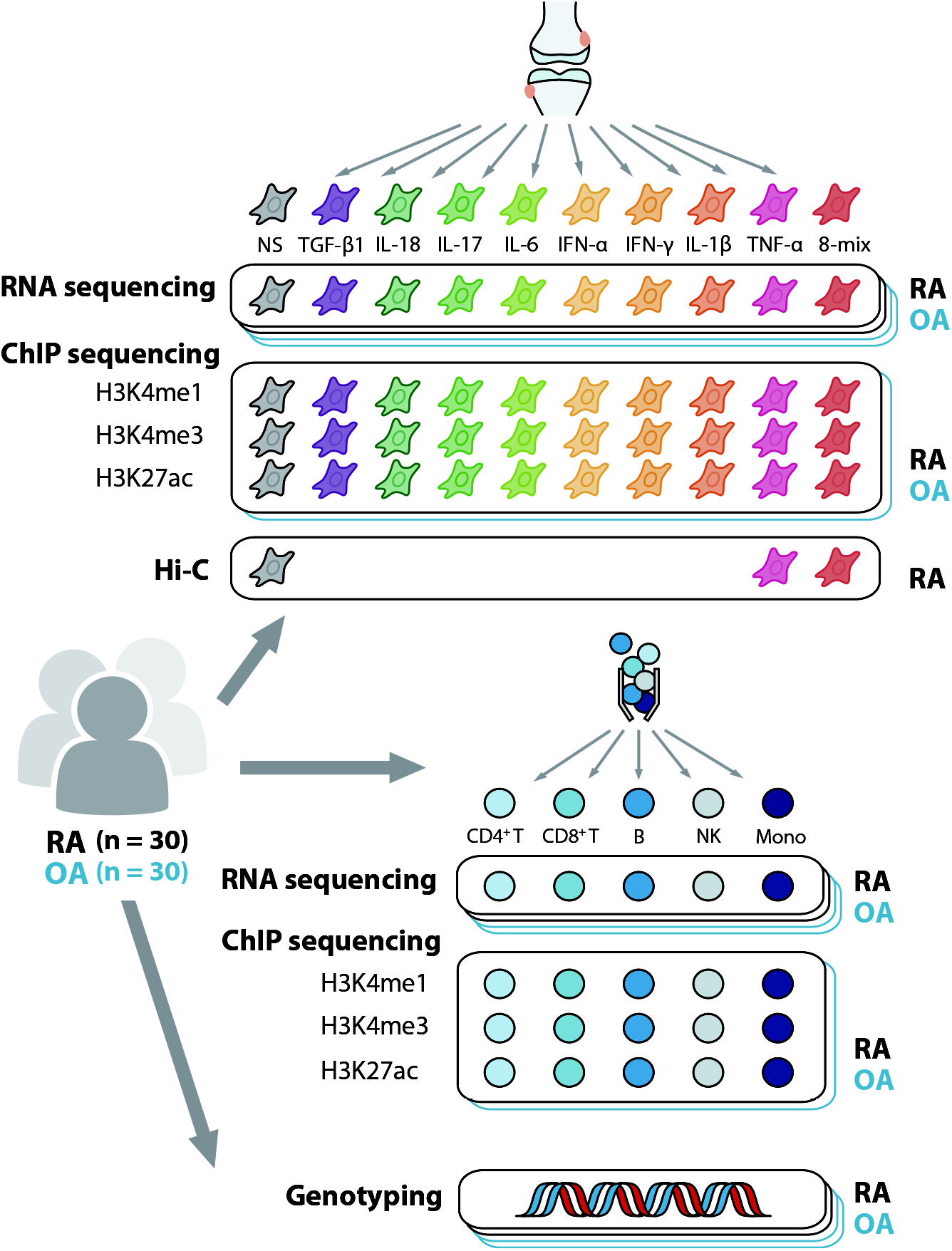
Experimental design for integrative analysis of activated SFs from RA and OA patients. Our study design included SFs stimulated by 8 different factors plus a combination of all the factors. Specifically, cells were treated for 24 h with one of the following: IFN-α 100 U/mL, IFN-γ 200 U/mL, TNF-α 10 ng/mL, IL-1β 10 ng/mL, IL-6/sIL-6R 200 ng/mL, IL-17 10 ng/mL, TGF-β1 10 ng/mL or IL-18 100 ng/mL or 8-mix, a mixture of the above 8 cytokines. In addition, we used 5 freshly isolated PBMC populations (CD4^+^ T cells, CD8^+^ T cells, B cells, NK cells, monocytes) from the same patient cohort. RNA sequencing of individual samples from RA and OA patients (n=30 per each) was carried out, and ChIP sequencing and Hi-C analysis were conducted with pooled samples. SNP genotyping array was performed in all patients. SFs, synovial fibroblasts; PBMCs, peripheral blood mononuclear cells; RA, rheumatoid arthritis; OA, osteoarthritis; NS, non-stimulated.

First, we compared transcriptomes of SFs between the 10 stimulatory conditions and the 2 diseases. Principal component analysis (PCA) showed that the 8-mix induced a distinct signature compared with the other stimulations, corresponding to the first component (PC1 in figure 2A,B). For instance, the markedly high expression of *IL6* (Primary component loading on PC1 = −0.76, PC2 = 0.26) and other pathogenic genes (e.g., *MMP3*, *CSF2*) was achieved by the 8-mix (figure 2C). On the other hand, RASFs showed different signatures from OASFs irrespective of the stimulations, corresponding largely to the second component (PC2 in figure 2A,B).

**Figure 2.**
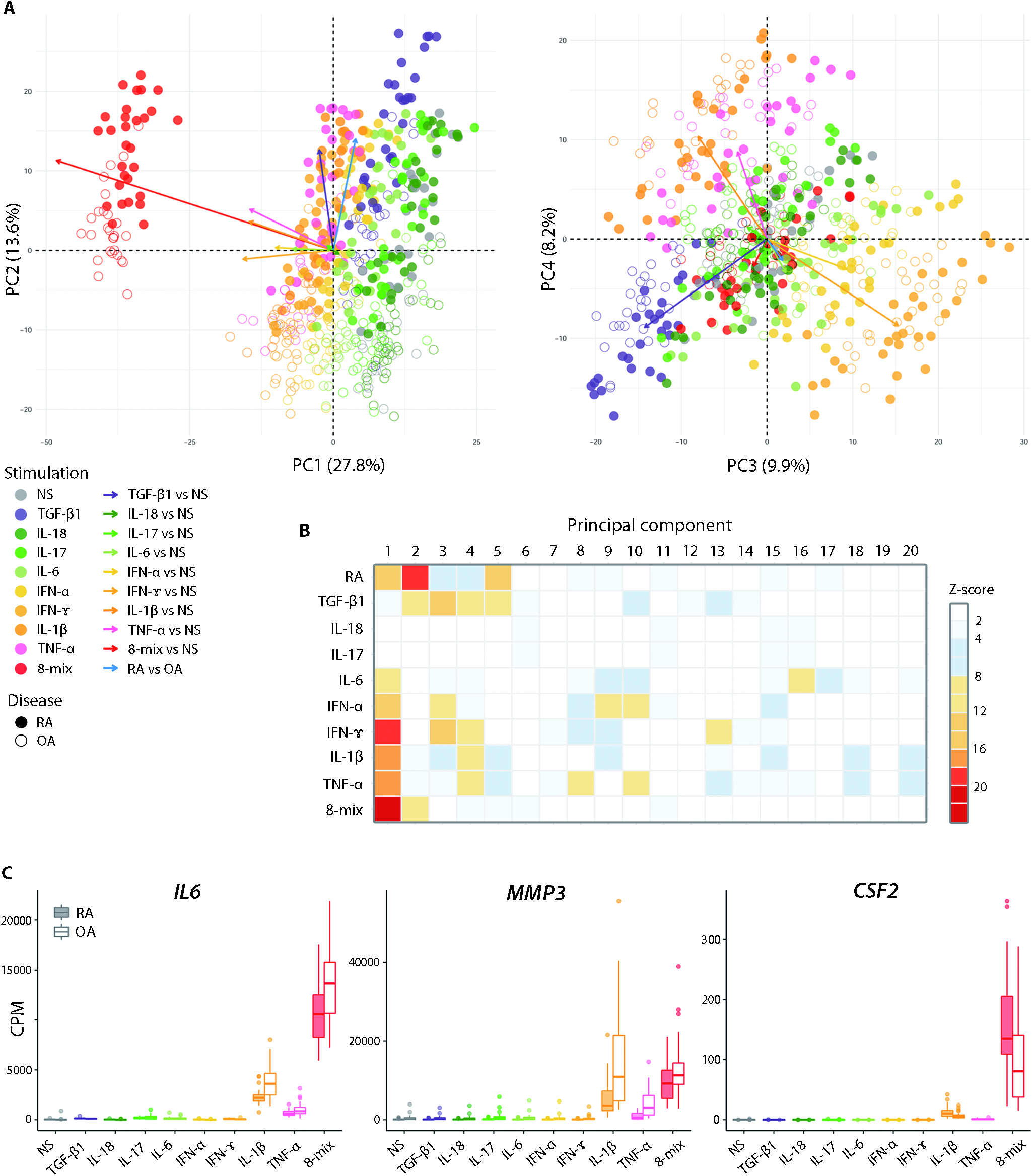
A distinctive transcriptome signature in SFs induced by the mixture of 8 cytokines. (**A**) Principal Component Analysis (PCA) of gene expression levels for the top 1000 variable genes. Samples projected onto PC1/PC2 (left) or PC3/PC4 (right). Numbers in parentheses indicate contribution ratio (percentage of variation) of the first 4 PCs. Arrows link the centroid of indicated groups and adjusted to start from the origin. (**B**) Summary of PCA for top 1000 variable genes. We fit a linear model to each PC (i.e., PC ~ Disease + Stimulation). Then we transformed the *P* values to normal Z-scores. (**C**)Transcript abundances of representative RA pathogenic genes (*IL6*, *MMP3* and *CSF2*) from RNA sequencing data in stimulated SFs. Boxes, interquartile range; whiskers, distribution; dots, outliers. PCA, principal component analysis; RA, rheumatoid arthritis; OA, osteoarthritis; NS, non-stimulated; CPM, count per million.

Recently, Fan Zhang and colleagues reported that SFs could be clustered into 4 fractions (F1-F4) by single-cell transcriptomics of synovial cells.[12] To elucidate the association between transcriptional changes of activated SFs and those 4 fractions, we scored our samples utilizing the reported gene sets that were characteristic of each of the fractions (online supplementary figure 1). [13] Interestingly, the F4 score, which was the population abundant in the lining of OA synovium,[12] was higher in OASFs irrespective of the stimulations. In contrast, the F2 score, which was a major *IL6* producer in the sublining, was strongly upregulated under IFN-γ, IFN-α, IL-6 or 8-mix stimulation. Accordingly, we surmised that certain SF populations were quantitatively stable between the diseases, and some are inducible under inflammation.

### eQTL effects of RA risk loci are modulated by an environmental perturbation

We next performed cis-eQTL analysis to evaluate the effect of genetic variants on gene expressions. Together with tissue-by-tissue analysis, we also utilized Meta-Tissue software (online supplementary materials) for a meta-analysis across SFs under 10 stimulations and 5 PBMC subsets. When we analyzed RA and OA samples separately, the eQTL effect sizes showed high similarities between the diseases. To improve analytical ability, we jointly analyzed the RA and OA samples. We considered variants with FDR < 0.1 in single tissue analysis or *m*-value > 0.9 in meta-analysis as significant eQTLs.[14, 15]

As a result, 3245 - 4118 genes in SFs and 2557 - 2828 genes in PBMCs had significant eQTLs. In total, 2368 genes showed eQTL effects only in SFs (online supplementary figure 2A,B). A tissue-wise difference of eQTL effects was indicated by a distinguishable pattern of eQTLs in SFs from that of PBMCs when comparing the effect sizes in meta-analysis (online supplementary figure 2C). When we focused on stimulated SFs, the 8-mix showed the smallest correlation coefficients compared with other conditions.

Furthermore, we examined candidate causal genes among RA risk loci in SFs. We focused on eQTL variants with minimum *P* values in each associated eGene with meta-analysis or single tissue analysis, which are in linkage disequilibrium (LD) with GWAS top-associated loci (online supplementary table 1). One example is rs6074022, which is in tight LD (r^2^ = 0.95 in EUR, r^2^ = 0.9 in EAS population) with an established RA risk SNP rs4810485.[16] rs6074022 had robust eQTL effects on *CD40* in SFs, especially under 8-mix or IFN-γ stimulations (figure 3A,D). Importantly, the presence of an active regulatory region at rs6074022 was inferred only under these conditions (figure 3B). Although *CD40* is also expressed by B cells (figure 3C), the eQTL effect was not observed at this locus (figure 3D).

**Figure 3.**
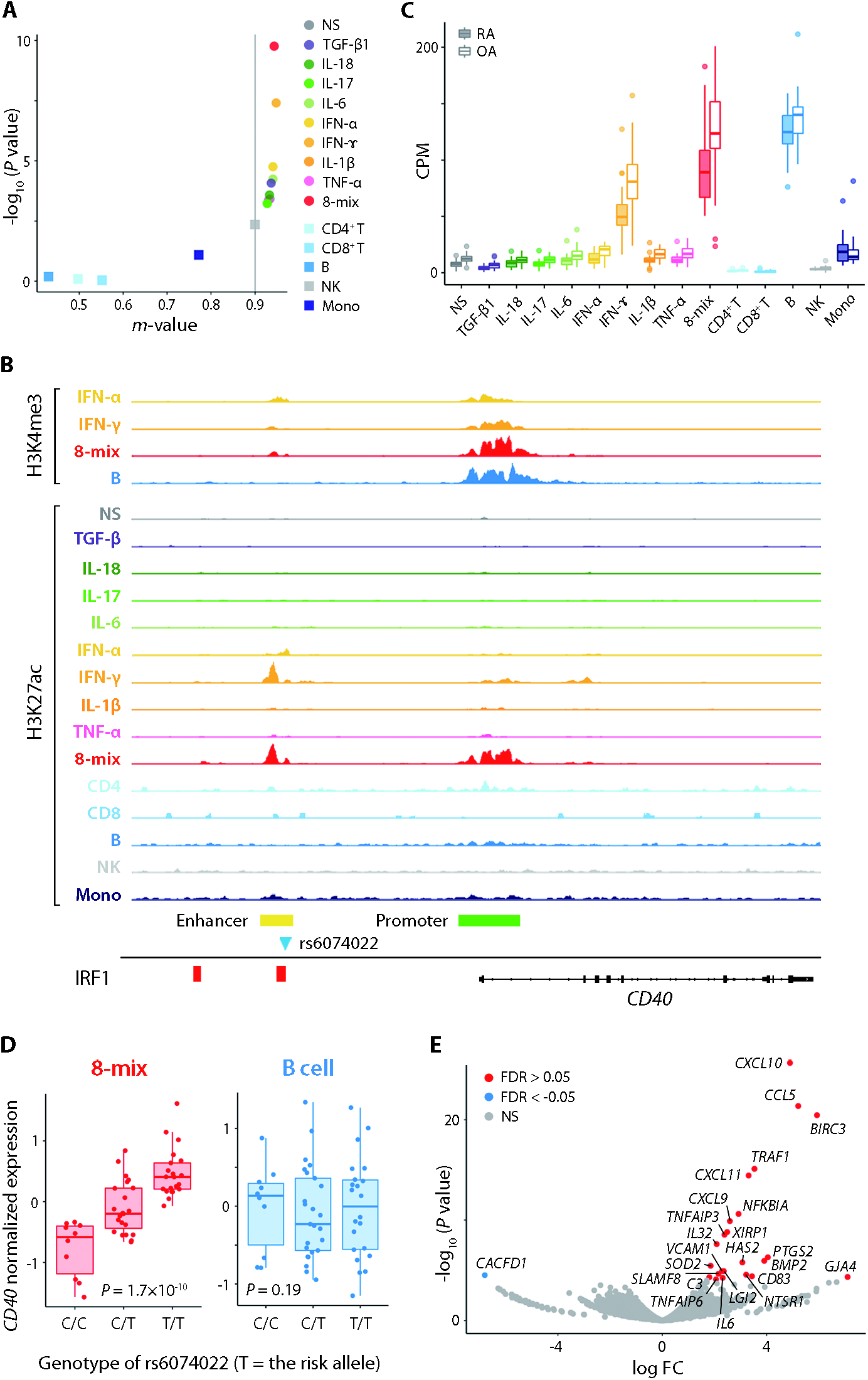
Stimulation-specific function of RA genetic risk loci. (**A**) dot plot of rs6074022-*CD40* cis-eQTL meta-analysis posterior probability *m*-values versus tissue-by-tissue analysis −log10 *P* value. The gray solid line (*m*-value = 0.9) corresponds to the significance threshold in this study. (**B**)Transcriptional regulatory regions around the *CD40* gene and positional relationship of rs6074022 (blue triangle) in stimulated SFs and PBMCs. IRF1 biding sites were obtained from the public epigenome browser ChIP-Atras. Data were visualized using the Integrative Genomics Viewer (IGV). (**C**) Transcript abundance of *CD40* from RNA sequencing data in stimulated SFs and PBMCs. (**D**) Expression of *CD40* in SFs stimulated by the mixture of 8 cytokines (left) and B cells (right) from each individual plotted according to the rs6074022 genotype. Nominal *P* values in eQTL mapping are shown. (**E**) A volcano plot of differential gene expression analysis comparing the presence or absence of CD40 ligand (CD40L) for IFN-γ-stimulated SFs. Orange and blue points mark the genes with significantly increased or decreased expression respectively for the addition of CD40L (FDR <0.01). Boxes, interquartile range; whiskers, distribution; dots, outliers in (**C**) and (**D**). RA, rheumatoid arthritis; OA, osteoarthritis; NS, non-stimulated; CPM, count per million.

The biological role of the CD40-CD40L pathway in SFs has been discussed.[17, 18] We performed transcriptomic analysis of RASFs stimulated with a 2-trimer form of the CD40 ligand and IFN-γ. As a result, some cytokines (e.g., *IL6*) and chemokines (e.g., *CCL5*, *CXCL10*) were significantly upregulated by the ligation of CD40L (figure 3E). Taken together, we conjecture that CD40 expression in SFs is influenced by genetic and environmental predisposition, and the CD40-CD40L pathway might have a pathogenic role in RASFs. Other examples of eQTL-eGene pairs in SFs are shown in online supplementary figure 3.

### Genetic risk of RA accumulate in the transcriptomic and epigenomic perturbations induced by synergistic proinflammatory cytokines

Next, in order to elucidate the link between RA genetic risk and transcriptomic perturbations of activated SFs, we performed gene-set enrichment analysis with RA risk loci using MAGMA software (online supplementary materials). This analysis indicated that perturbed gene sets subject to IFN-α, IFN-γ and 8-mix stimulation significantly overlapped with RA risk loci (*P* = 2.0 × 10^−4^, 1.7 × 10^−4^ and 8.3 × 10^−6^, respectively) (online supplementary figure 4A,B). Those data contrasted with the non-significant association of transcriptome differences between the diseases with RA genetic risk. These findings indicate that there is an accumulation of RA genetic risk in the pathways that are perturbed under specific conditions in SFs.

We also assessed the enrichment of GWAS top-associated loci in regulatory regions including super-enhancers (SEs). SEs are large clusters of enhancers collectively bound by an array of transcription factors (TFs) to define cell identity, and are hotspots for disease susceptibility.[19–21] Although the significant overlap of SEs in Th cells and B cells with RA risk loci has been reported,[19] SFs have not been examined. Here we compared the enrichment of RA risk loci to SEs and typical-enhancers (TEs), and risk loci showed significant enrichment with SEs in CD4^+^ T cells and B cells, as well as with SEs in 8-mix stimulated SFs (figure 4A). Some risk loci showed overlap with SEs which appear uniquely under 8 mix treatment (figure 4B,C). When we performed a similar analysis using risk loci for type 1 diabetes mellitus (a representative non-articular autoimmune disease), only SEs in CD4^+^ T cells and B cells showed significant enrichment (online supplementary figure 4C). The number or width of 8-mix SEs were comparable to those in other stimulations (online supplementary figure 5). Consequently, SFs might behave as key players in RA pathogenesis especially under synergistic inflammation.

**Figure 4.**
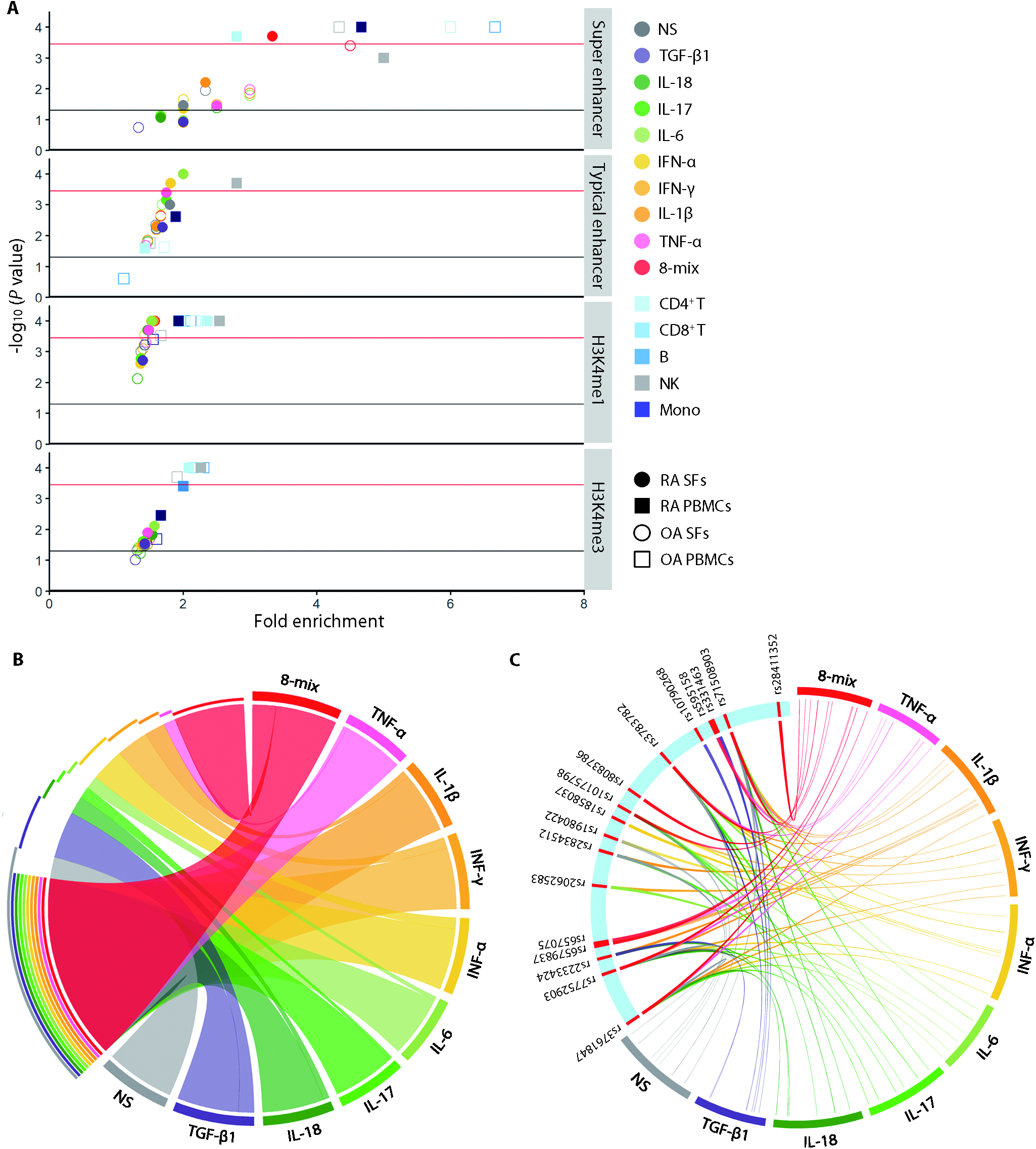
Enrichment of RA genetic risk in SFs super-enhancers under 8 cytokine stimulation. (**A**) Enrichment of RA risk loci in transcriptional regulatory regions of stimulated SFs and PBMCs. Active enhancers were classified into super-enhancers (SEs) and typical-enhancers (TEs) following standard ROSE algorithms. The red solid lines (−log10 (*P* value) = 3.2) and the black solid lines (−log_10_ (*P* value) = 1.3) are the cutoffs for Bonferroni significance and nominal *P* = 0.05, respectively. (**B**) A circus plot showing the overlap of SEs in SFs under different stimulatory conditions. Only the regions unique to each condition or common to all of the conditions are depicted. (**C**) A circus plot showing the overlap of RA risk loci and SEs in SFs under different stimulatory conditions. NS, non-stimulated; SE, super-enhancer; TE, typical enhancer.

### SEs induced by cytokine mixtures regulate genes crucial for RA pathogenesis

Following the results above, we attempted to elucidate the genes regulated by 8-mix-enhanced SEs. First, we combined the 3D genome architectures (chromatin loops detected by Hi-C analysis), the position of SEs, promoter regions (defined with H3K4me3 ChIP sequencing analysis) and genomic coordinates. We annotated “SE-contacted genes” such that one side of Hi-C loop anchors overlapped a SE, the other side coincided with the TSS and coexisted with the H3K4me3 peak (figure 5A). SEs were highly overlapped with Hi-C loop anchors than were TEs or H3K4me1 peaks (online supplementary figure 6A). When the TSS and H3K27ac peak was connected by a Hi-C loop, the variation of mRNA expression showed a significant correlation with the H3K27ac peak variation (online supplementary figure 6B). These results underscore the validity of connecting active enhancer marks and TSS by Hi-C loops as previous reports.[22, 23] When we compared the expression of SE-contacted genes and TE-contacted genes, the former showed significantly higher expression than the latter (online supplementary figure 6C).[21]

**Figure 5.**
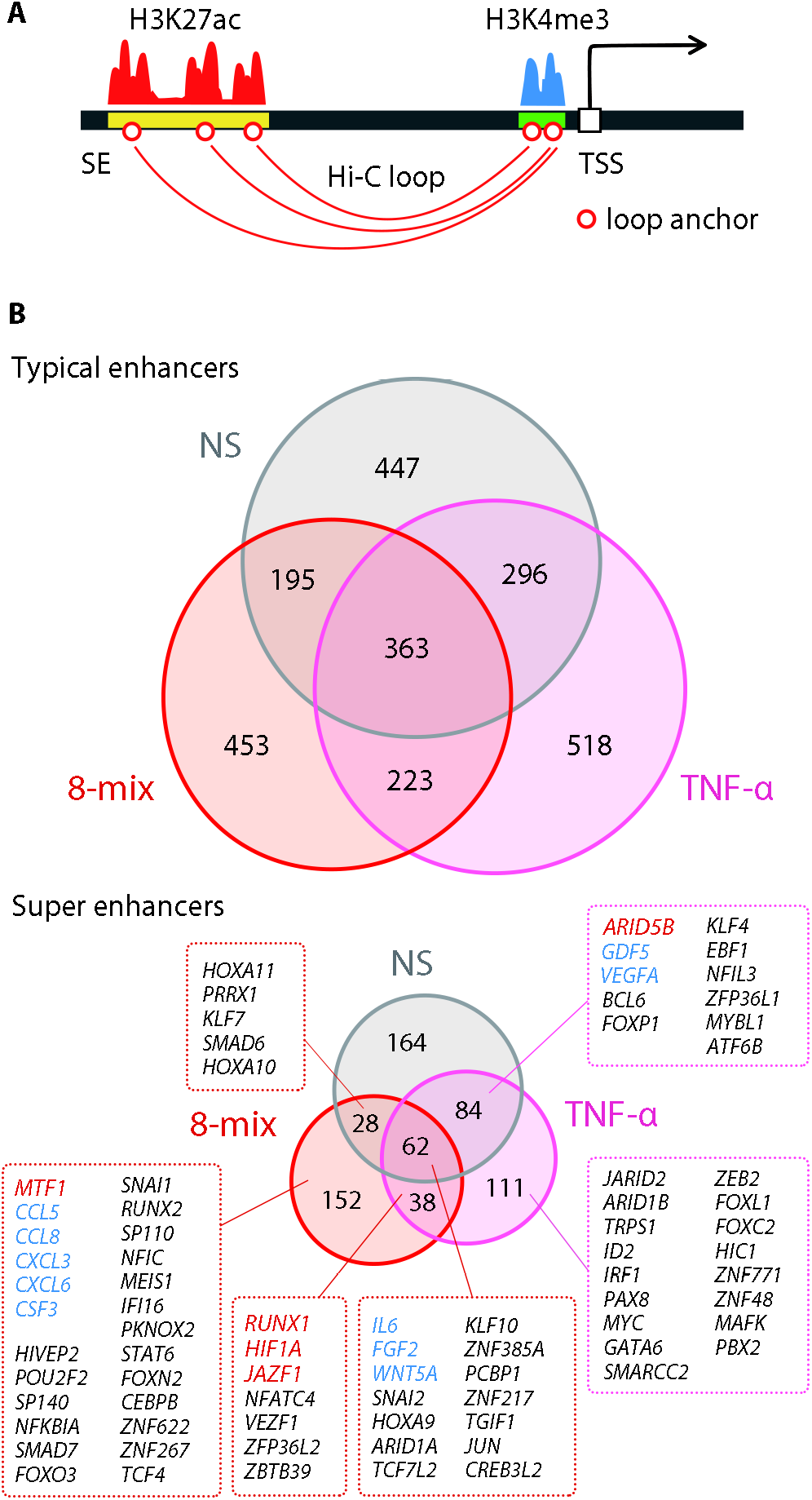
RA pathogenic factors regulated by SEs in SFs treated with 8 cytokines. (**A**) A schematic image of “SE-contacted genes”. (**B**) A Venn diagram representing the overlap of TE-contacted (top) or SE-contacted (bottom) genes in SFs under different stimulatory conditions. Red, blue and black text highlight genes whose contacted SEs overlap with RA risk loci, cytokines and chemokines and transcription factors, respectively. TSS, transcriptional start site; NS, non-stimulated; SE, super-enhancer; TE, typical enhancer.

Next, we compared genes contacted by either SEs or TEs of SFs under 3 different conditions: non-stimulated, TNF-α or the 8-mix. The proportions of overlap between these conditions were smaller in SE-contacted genes (9.7%) than TE-contacted genes (14.5%) (figure 5B), indicating that the stimulation-specific expression profile and SEs formation. SE-contacted genes included a number of TFs (e.g., *MTF1, RUNX1*), cytokines (e.g., IL6) and chemokines (e.g., *CCL5, CCL8*) (figure 5B, online supplementary table 2).

*IL6* is a representative example of an 8-mix SE-contacted gene. Although this gene is regulated by a SE (almost 30 kb long) that exists upstream of the TSS in non-stimulated or TNF-α stimulated SFs, this SE elongates to 70 kb long under the 8-mix and an additional Hi-C loop emerges (figure 6A). When we inhibited SE formation with JQ1, a BRD4 inhibitor, the increased IL-6 expression under the 8-mix was disturbed (figure 6B). The necessity of SEs for elevated IL-6 production under synergistic inflammation was inferred.

**Figure 6.**
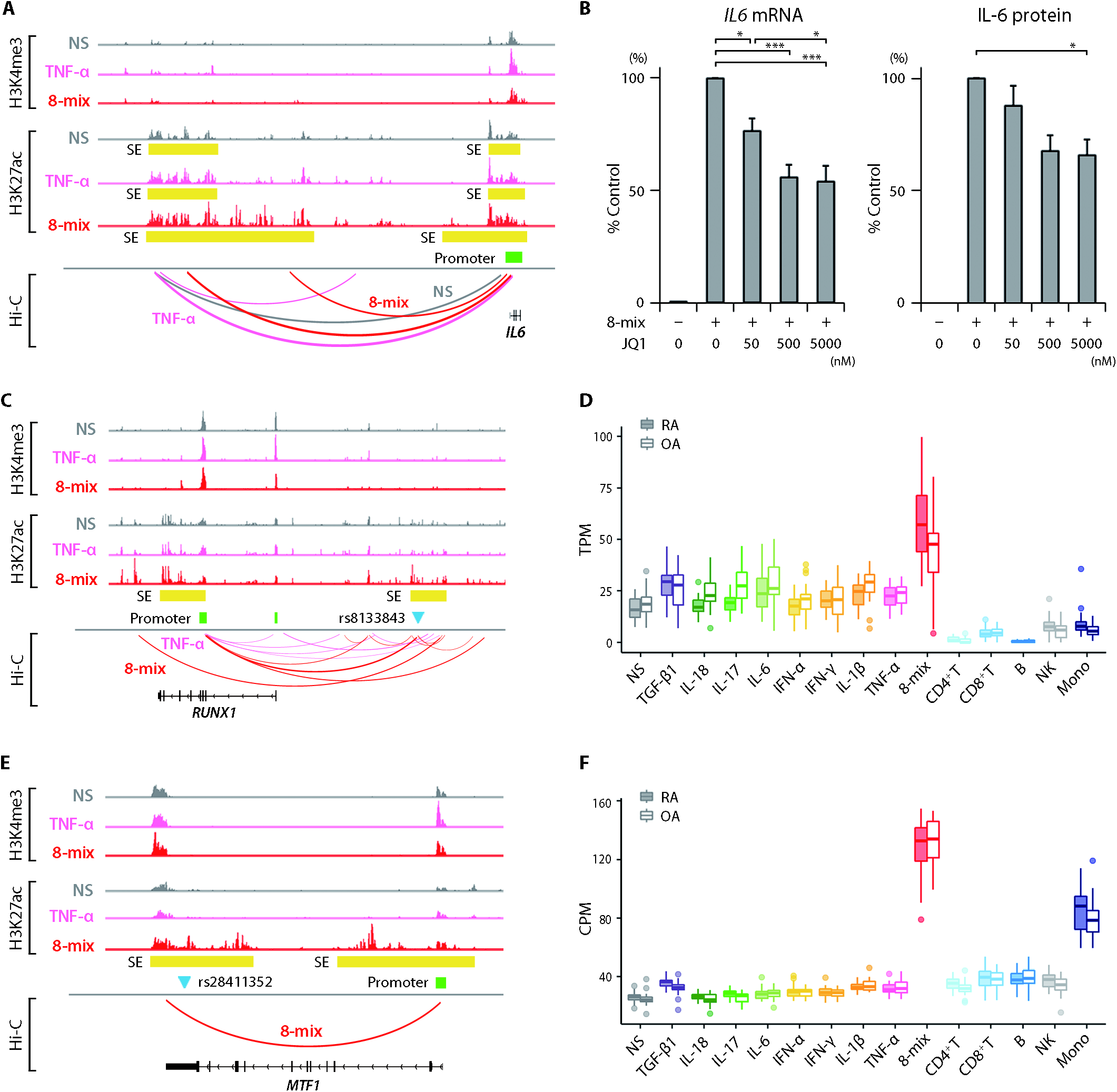
Representative SE-contacted genes in SFs treated with 8 cytokines. (**A, C, E**) Organization of transcriptional regulatory regions around *IL6* (**A**), *RUNX1* (**C**) and *MTF1* (**E**) genes and positional relationship of RA risk loci (blue triangle, rs8133848 for *RUNX1* and rs28411352 for *MTF1*) and chromatin conformation in stimulated SFs. Data were visualized using the Integrative Genomics Viewer (IGV). (**B**) Expression of IL-6 in SFs treated with JQ1. The mRNA and protein expression were quantified by qRT-PCR (left) and ELISA (right), respectively. Bar graphs, mean; Error bars, SEM. *P* values were determined using one-way ANOVA followed by Tukey’s multiple comparison test (*, *P* < 0.05, ***, *P* < 0.001). (**D, F**) Transcript abundances of *RUNX1* isoform (RUNX1b) (**D**) and *MTF1* (**F**) from RNA sequencing data for stimulated SFs and PBMCs. Boxes, interquartile range; whiskers, distribution; dots, outliers. RA, rheumatoid arthritis; OA, osteoarthritis; NS, non-stimulated; TPM, transcripts per million; CPM, count per million; SE, super enhancer.

Another example is *Runt related transcription factor 1* (*RUNX1*), a master-regulator involved in normal and malignant hematopoiesis.[24–26] The binding motif of *RUNX1* was enriched in SEs of synovial fluid-derived CD4^+^ T cells from juvenile idiopathic arthritis patients.[27] In our study, RA risk locus rs8133843 overlapped with an 8-mix SE that exists upstream of *RUNX1* (figure 6C). A Hi-C loop was formed with the promoter immediately above the second exon of *RUNX1* only in the 8-mix, and the *RUNX1* expression was higher in the 8-mix compared with others (figure 6D).

*Metal-regulatory transcription factor-1* (*MTF1*), a zinc finger TFs, is another example. RA risk locus rs28411352 overlapped with an 8-mix unique SE that exists upstream of the *MTF1* (figure 6E). The Hi-C loop was detected with the promoter only under the 8-mix. *MTF1* expression was upregulated in the 8-mix (figure 6F).

### Transcription factors associated with stimulation-induced SE formation control arthritis progression

Finally, we searched for candidate modulators that were crucial for SE formation, especially in the 8-mix. In the previous study, key TFs for SE formation were reported to be controlled by SEs themselves, forming a self-regulatory network.[21] From this perspective, we used motif analysis to focus on SE-contacted TFs that were also enriched in 8-mix SEs and compared them with TEs or unstimulated SEs (figure 7A). Among SE-contacted genes, TFs such as SNAI1, TCF4 and MTF1 showed significant motif enrichment in 8-mix SEs. MTF1 was the only example that also showed the overlap of 8-mix SE and RA risk variant (figure 6E). Although the RUNX1 motif was not enriched in 8-mix SEs compared with unstimulated SEs, its motif was significantly enriched in SEs compared with the background sequence, both in 8-mix and without stimulation (*P* = 1.0 × 10^−16^ and 1.0 × 10^−20^, respectively). *In vitro* validation analysis showed that the expression of 8-mix SE-contacted genes was significantly suppressed by *MTF1* and *RUNX1* knockdown (*P* = 2.0 × 10^−3^ and 4.3 × 10^−4^, respectively) (figure 7B, online supplementary figure 7). The effect of *MTF1* knockdown was more pronounced in 8-mix SE-contacted genes than TE-contacted genes. In *in vitro* assay, the increased 8-mix SE-contacted gene expressions (IL-6, CCL5) from stimulated RASFs was reduced by treatment with APTO-253, a MTF1 inhibitor (figure 7C,D).[28] Furthermore, in a collagen induced arthritis (CIA) model, APTO-253 demonstrated significant preventive (online supplementary figure 8) and therapeutic activity (figure 7E,F) on arthritis formation. Collectively, these results indicated that certain TFs play critical roles in the formation of epigenomic structures induced by synergistic proinflammatory cytokines, and support MTF1 inhibition as a promising therapy candidate for RA (online supplementary figure 9).

**Figure 7.**
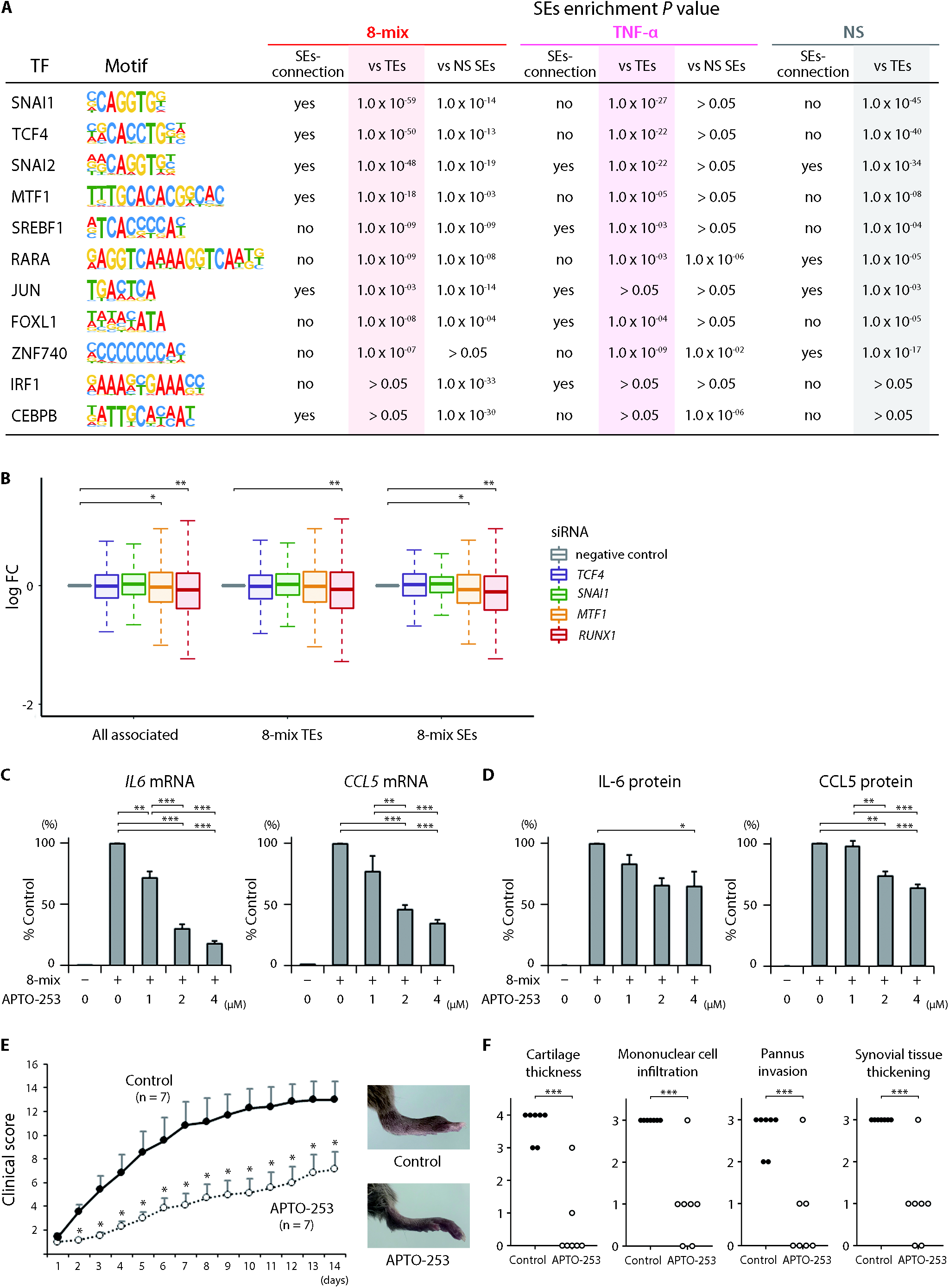
Transcription factors associated with stimulation-induced SE formation and arthritis progression. (**A**) Table depicts transcription factor binding motifs enriched at SEs in stimulated SFs. Following are summarized: attribution to SE-contacted genes, relative enrichment *P* values to TEs in each stimulatory condition or to SEs of non-stimulated condition. (**B**) Expression of SE- or TE-contacted genes in SFs stimulated by 8 cytokines in cells depleted of specified transcription factors (*TCF4*, *SNAI1*, *MTF1* and *RUNX1*) relative to control SFs. Boxes, interquartile range; whiskers, distribution. *P* values, paired t test (*, *P* < 0.05, ***, *P* < 0.001). (**C, D**) Expression of IL-6 and CCL5 in SFs treated with APTO-253. The mRNA and protein level were quantified by qRT-PCR (**C**) and ELISA (**D**), respectively. Bar graphs, mean; Error bars, SEM. *P* values, one-way ANOVA followed by Tukey’s multiple comparison test (*, *P* < 0.05, **, *P* < 0.01, ***, *P* < 0.001). (**E, F**) Therapeutic effect of APTO-253 on CIA model. Following the onset of arthritis, mice were intravenously injected with either control or 15 mg/kg APTO-253 for twice per day for 2 consecutive days per week. Clinical (**E**) and pathological scores (**F**). Dots, mean; Error bars, SEM. *P* values, Mann–Whitney U test (*, *P* < 0.05, ***, *P* < 0.001). NS, non-stimulated; SE, super enhancer; TE, typical enhancer.

## Discussion

In this study, we were able to describe the dynamic landscape of activated SFs and their contribution to RA pathogenesis. Recent large-scale eQTL studies enhanced our understanding of complex diseases.[29, 30] Considering the importance of studying disease-relevant tissues for functional understanding of GWAS variants,[14, 31] we conducted cis-eQTL analysis of SFs, which are major local effector cells in arthritic joints. One example of eQTL-eGene pairs in SFs was the association of an RA risk SNP (rs4810485) with *CD40* expression. Although no significant eQTL effects of this locus in B cells was observed in our data, a previous report showed the protein-level QTL association of CD40 in CD19^+^ B cells and this locus in Europeans.[32] On the contrary, a larger eQTL study by the GTEx consortium that showed genome-wide significant eQTL effects of rs6074022 on CD40 in lung (mainly fibroblasts) (*P* = 3.1 × 10^−26^) and cultured fibroblasts (*P* = 8.4 × 10^−16^), but not in EBV-transformed lymphocytes (*P* = 0.04).[33] The subtle gap in results could be derived from racial differences of study populations and study design. Naturally, although the possible importance of CD40 signals in B cells for RA pathogenesis cannot be neglected, our result shed light on the role of CD40-CD40L signals in a genetic network of synovitis.

The significance of synergistic interactions between proinflammatory cytokines and SFs is supported by the observed accumulation of RA risk loci as shown by transcriptomic and epigenomic data. The GWAS-SEs enrichment analysis shows that SFs can behave as pathogenic cells in the development of RA (analogous to CD4^+^ T cells and B cells) especially when exposed to the synergistically acting cytokines. Furthermore, recent fine chromatin contact maps revealed that SEs are in close proximity to the promoter of the gene they activate.[34–37] During cytokine synergy, Hi-C analysis suggested that there were dynamic conformational changes in three-dimensional structures involving SEs and the promoter of pathological molecules. We found marked expression of 8-mix stimulated SE-contacted genes (i.e., *IL6*, *RUNX1*, *MTF1*).

SEs can collapse when their co-factors (e.g., BET family) are perturbed.[38] In RASFs, BRD4 silencing or inhibitor reduced cytokine secretion, migration and invasion activity, and ameliorated experimental arthritis.[39] On the other hand, selective modulation of disease-associated SEs in a cell type-specific manner may have better safety profiles than pan-SEs inhibitors (i.e., JQ1). In T cell leukemias, a small monoallelic insertion creates binding motifs for the master regulator MYB, which trigger SEs initiation upstream of oncogenes.[25] In the present study, we analyzed TFs that have the potential to be selective SE modulators in activated SFs. Our results suggested that MTF1 participates in SE formation, putatively making a feedback loop to maintain the epigenomic machinery. MTF1 regulates gene expression in response to zinc and various stresses.[40] In the setting of disease, MTF1 could contribute to tumor metastasis and chemoresistance in some cancer cells.[41] APTO-253, a MTF1 inhibitor,[28] is a small molecule with antitumor activity against a wide range of human malignancies in *in vitro* and *in vivo*.[42, 43] In the present study, APTO-253 demonstrated inhibitory activity on the expression of pathogenic molecules (IL-6, CCL5) from RASFs and antiarthritic activity in a CIA model. Previously, the zinc-ZIP8-MTF1 axis was identified as a catabolic regulator of cartilage destruction.[44] Furthermore, intra-articular injection of adenovirus expressing-MTF1 in an experimental mouse model of OA promoted the expression of various matrix-degrading enzymes, cytokines and chemokines in SFs. This evidence supports the essential role of MTF1 as a genetic hub for joint destruction and inflammation.

There are some limitations of this study. First, the number of patients included in the cis-eQTL analysis was limited owing to sample accessibility, resulting in putatively large false negatives. Second, SFs were purified by serial passages. Previous reports showed that passage number could affect the epigenome and transcriptome of SFs.[45, 46] To minimize the impact of culture, we restricted our analysis to early passage cells. Third, isolated PBMCs were not artificially stimulated with cytokines *ex vivo*. Thus, it is not clear whether the eQTL difference between SFs and PBMCs is attributable to cell type difference. However, over 80% of RA patients had moderate to high disease activity at the time of blood sampling, indicating that PBMCs were subject to a proinflammatory environment.

Overall, our findings shed light on the concept of SF-targeted therapy from the perspective of epigenome remodeling. Some TFs (MTF1, RUNX1) preferentially recruited to SEs during exposure to synergistic proinflammatory cytokines would constitute novel drug targets.

## Acknowledgments

We would like to thank G Inoue, M Abe, K Myouzen and K Kobayashi for their technical assistance at the Laboratory for Autoimmune Diseases, and Dr. Y Momozawa and Dr. M Kubo for providing technical advice at the Laboratory for Genotyping Development, RIKEN. We received generous support from all the physicians who participated in sample collection at the Department of Orthopaedic Surgery, the University of Tokyo.

## Funding

This research was supported by funding from Takeda Pharmaceutical Co., Ltd. (Y. Kochi., K. Y. and K.F.), the Ministry of Health, Labour and Welfare, Ministry of Education, Culture, Sports, Science and Technology KAKENHI Grant-in-Aid for Scientific Research (B) (18H02846) and Grant-in-Aid for Scientific Research (C) (17K09972) from the Japan Society for the Promotion of Science.

## Author contributions

H.T., S.S., K.I., Y. Kochi., K.Y. and K.F. designed the research project. M.O. conducted bioinformatics analysis on the advice of K.I.. A.S. performed RNA sequencing. H.T. performed ChIP sequencing and other in vitro experiments. T.S. and K.S. performed Hi-C. H.T., Y.T., H.I., J.H., Y. Kadono. and S.T. contributed human samples. S.S. performed figure editing. H.T. and M.O. wrote the manuscript with critical inputs from Y. Kochi., K.Y. and K.F..

## Competing interests

None declared.

## Data and materials availability

The datasets generated during this study are available at the National Bioscience Database Center (NBDC) with the study accession code hum0207 (read counts data of RNA sequencing, hum0207.v1.RNA.v1; eQTL summary, hum0207.v1.eQTL.v1; peaks data of ChIP sequencing, hum0207.v1.ChIP.v1; chromatin loops data of Hi-C, hum0207.v1.HiC.v1).

## Notes

### Competing Interest Statement

The authors have declared no competing interest.

### Summary of Updates

Main document updated; Supplementary file updated.

