## Supplementary File for "Parsing multiomics landscape of activated synovial fibroblasts highlights drug targets linked to genetic risk of rheumatoid arthritis"

**Online supplementary materials and methods**

**Study Design**

The overall objectives were to explore the pathogenic mechanisms of RASFs through elucidating the genetic contribution to molecular regulatory networks under inflammatory condition. First, we quantified mRNA expression by RNA sequencing and compared transcriptome of SFs between the 10 conditions (i.e., non-stimulated, IFN-α, IFN-γ, TNF-α, IL-1β, IL-6/sIL-6R, IL-17, TGF-β1, IL-18 or 8-mix) and the diseases (i.e., RA, OA). We next performed cis-eQTL analysis to evaluate the effect of genetic variants on gene expressions in stimulated SFs and five major immune cell subsets ($CD4^+$ T cells, $CD8^+$ T cells, B cells, NK cells, monocytes) from the same patient. In addition, we examined candidate causal genes among RA risk loci in SFs. Focusing on eQTL variants in LD with GWAS top-associated loci, we assessed the biological role of the CD40-CD40L pathway in SFs by transcriptomic analysis of RASFs stimulated with a 2-trimer form of the CD40 ligand and IFN-γ as a representative example. Next, in order to elucidate the link between RA genetic risk and transcriptomic and epigenomic perturbations of stimulated SFs, we performed a gene-set enrichment analysis with RA-associated genetic loci, and assessed the enrichment of GWAS top-associated loci in regulatory regions (SEs or TEs) identified with ChIP sequencing. Furthermore, we combined the 3D genome architectures (chromatin loops detected by Hi-C analysis), the position of SEs, promoter regions (defined with H3K4me3 ChIP sequencing analysis) and genomic coordinates in SFs under 3 different conditions: non-stimulated, TNF-α or the 8-mix. To reveal candidate modulators crucial for SE formation, especially in the 8-mix, we integrated motif analysis to focus on SE-contacted TFs that were also enriched in 8-mix SEs and compared them with TEs or unstimulated SEs. Finally, the promising pathogenic TFs (i.e., MTF1, RUNX1) in RASFs were silenced with siRNAs to validate transcriptomic effects, and the impact of MTF1 inhibitor was assessed by *in vitro* and *in vivo* assay.

**Patient and public involvement**

Patients and/or the public were not involved in the design, or conduct, or reporting, or dissemination plans of this research.

**Human Participants and Sample Collection**

Synovial tissues were obtained from RA and OA patients (n = 30 each) undergoing joint replacement surgery at the University of Tokyo Hospital, Japan. RA patients fulfilled the 2010 ACR/EULAR (American College of Rheumatology/European League Against Rheumatism) criteria for the classification of RA.[1] Patient characteristics are summarized in online supplementary table 4. This study was approved by the Ethics Committees of the University of Tokyo (G3582), RIKEN and the indicated medical institutions. Written informed consent was obtained from each subject in accordance with the Declaration of Helsinki. Fresh synovial tissues were minced and digested with 0.1% collagenase (Worthington) at 37°C, in 5% $CO_2$ for 1.5 h and SFs were cultured in Dulbecco's modified Eagle's medium (DMEM; SIGMA) supplemented with 10% fetal bovine serum (FBS; BioWest), 100 µg/mL L-glutamine, 100 U/mL penicillin, 100 µg/mL streptomycin (all from Invitrogen). SFs from passage 2 or 3 were used for RNA sequencing, ChIP sequencing, Hi-C and functional studies after removal of macrophages by magnetic separation with CD14 microbeads (Miltenyi Biotec). The purity of SFs was tested by flow cytometry analysis (MoFlo XDP; Beckman

Coulter). SFs were stained with CD14-, Thy-1 (CD90)-specific monoclonal antibodies (clone IDs: M5E2, 5E10, respectively, all from BioLegend). Most cells (>99%) had the surface marker for fibroblasts (Thy-1) but not CD14.

We collected peripheral blood from the same patients. PBMCs were isolated using Ficoll-Paque density gradient centrifugation followed by staining with CD3-, CD4-, CD8-, CD14-, CD19-, and CD56-specific monoclonal antibodies (clone IDs: UCHT1, OKT4, RPA-T8, M5E2, HIB19 and HCD56, respectively, all from BioLegend). Five immune cell populations were sorted by flow cytometry (MoFlo XDP; Beckman Coulter) using the following gating strategy: $CD4^+$ T cells: $CD3^+CD4^+CD8^-CD19^-$; $CD8^+$ T cells: $CD3^+CD4^-CD8^+CD19^-$; B cells: $CD3^-CD19^+$; NK cells: $CD3^-CD14^-CD19^-CD56^+$; and monocytes: $CD3^-CD14^+CD19^-$. There were $3 \times 10^5$ cells in each population.

**RNA sequencing**

Purified SFs ($2 \times 10^4$ cells/well) were seeded with DMEM (10% FBS, 100 µg/mL L-glutamine, 100 U/mL penicillin, 100 µg/mL streptomycin) into a 24-well flat-bottom plate (Corning) and incubated at 37°C, in 5% $CO_2$. After 12 h, one of the following was added: 100 U/mL IFN-α (HumanZyme), 200 U/mL IFN-γ, 10 ng/mL TNF-α, 10 ng/mL IL-1β, 200 ng/mL IL-6/sIL-6R, 10 ng/mL IL-17 (all from PeproTech), 10 ng/mL TGF-β1 (R&D), 100 ng/mL IL-18 (MBL). Alternatively, cells were treated with “8-mix” (a mixture of the above 8 cytokines that simulated synergistic inflammation in arthritic joints). These 8 cytokines were selected on the basis of 1) existing therapeutic targets or 2) the number of articles that reported the cytokine as pathogenic on a public database (PubMed). The cells were stimulated for an additional 24 h at 37°C, in 5% $CO_2$.

Total RNA from SFs and freshly sorted PBMCs was isolated using AllPrep DNA/RNA/miRNA Universal Kit (Qiagen). Libraries for RNA sequencing were prepared using TruSeq Stranded mRNA Library Prep Kit (Illumina). RNA sequencing was carried out on Illumina HiSeq 2500 (read length of 125 bp, paired end).

**Bioinformatic analysis of RNA sequencing data**

RNA sequencing reads were aligned to the human genome assembly hg19/GRCh37 excluding minor haplotypes, random and unknown sequences. Alignment of the reads was performed by STAR (version 2.5.3) (Key resources information) based on the GENCODE v27 (GRCh37 version) annotation. We only utilized reads that were uniquely mapped (corresponding to a mapping quality of 255 for BAM files), and properly paired for further analysis.

Gene-level read counts were quantified with HTSeq (version 0.6.0) (Key resources information) based on the GENCODE v27, with strand-specific assay mode and the other default parameters. Transcript-level quantifications were calculated with RSEM (version 1.3.0) (Key resources information).

We assessed the quality of each RNA sequencing sample by calculating the mean expression correlation coefficient with other samples with the same stimulatory conditions, equivalent to D statistics as described elsewhere.[2] All samples satisfied more than 10 million uniquely mapped read counts. We excluded samples the D statistics of which were lower than 0.9, resulting in 29 excluded samples (4 non-stimulated, 4 in TNF-α, 3 in IFN-α, 2 in IFN-γ, 6 in IL-1β, 4 in TGF-β1, 2 in IL-17, 3 in 8-mix and 1 CD8$^{+}$ T cell). The remaining 856 samples were utilized for the analysis.

Differential expression analysis was performed with the edgeR package (Key resources information) with gene-level count data. For each comparison, genes whose expression was less than 10 in more than 90% of samples were excluded. Gender was included as a covariate for all the comparative analyses. RUVseq R package (Key resources information) was utilized for finding hidden factors using 1000 nonvariable genes with conditions as negative control genes and unstimulated samples as negative control samples. Gender and 3 RUV factors were considered covariates in differential expression analysis. Genes with FDR less than 0.05 in the glml RT test implemented in edgeR were regarded as differentially expressed genes.

MAGMA software (Key resources information) was applied for gene-set analysis of GWAS data. We utilized RA GWAS summary statistics of European ancestry and performed gene set enrichment analysis following the instruction by the authors.[3] Briefly, we carried out “gene analysis” for GWAS summary statistics using the 1000 genomes European panel for LD calculation and GENCODE v27 for gene annotation. Then we carried out “gene-set analysis” using log-fold change of gene expression between conditions as gene covariate and performed enrichment analysis.

**ChIP sequencing**

Purified SFs ($1 \times 10^5$ cells/well) were seeded with DMEM (10% FBS, 100 µg/mL L-glutamine, 100 U/mL penicillin, 100 µg/mL streptomycin) into 6-well flat-bottom plates (Corning) and incubated at 37°C, in 5% $CO_2$. After 12 h, 100 U/mL IFN-α, 200 U/mL IFN-γ, 10 ng/mL TNF-α, 10 ng/mL IL-1β, 200 ng/mL IL-6/sIL-6R, 10 ng/mL IL-17, 10 ng/mL TGF-β1, 100 ng/mL IL-18 or 8-mix was added. The cells were stimulated for additional 24 h at 37°C, in 5%

$CO_2$.

Pooled SFs and freshly sorted PBMCs from RA or OA patients (n = 20 each) were cross-linked with 1% formaldehyde for 15 min at room temperature. Chromatin was prepared from pellets of SFs ($1 \times 10^7$ cells) and PBMCs ($2 \times 10^7$ cells) using a CHIP-IT High Sensitivity Kit and CHIP-IT PBMC Kit (both from Active Motif), respectively. Sonication was carried out by Covaris S2 (Covaris). The shearing efficiency was analyzed by agarose gel electrophoresis after RNase treatment, reversion of crosslinking and purification of DNA. Sheared chromatin (3 µg) was immunoprecipitated using 4 µL of each rabbit polyclonal antibody (H3K4me1, H3K4me3, H3K27ac, all from Active Motif). Sheared chromatin was used as the input DNA. Immunoprecipitated DNA was quantified with the Qubit dsDNA HS Kit (Invitrogen). Libraries for ChIP sequencing were prepared using TruSeq ChIP Library Prep Kit (Illumina) with 5 ng of DNA fragments. DNA size selection (250 - 300 bp) was carried out by BluePippin (Sage Science). ChIP sequencing was carried out on an Illumina HiSeq 2500 (read length of 50 bp, single end).

**Bioinformatics analysis of ChIP sequencing data**

Sequencing reads from each ChIP sequencing sample were mapped to human genome assembly hg19/GRCh37 using Bowtie2 (Key resources information). PCR duplicates were removed, and only uniquely mapped reads were used for peak calling. MACS 2.0 (Key resources information) was used to detect peaks which were enriched in immunoprecipitated samples over the input. Peak calling was performed with narrow peak mode for H3K4me3 and H3K27ac and broad peak mode for H3K4me1.

NSC and RSC were calculated by cross-correlation analysis following ENCODE guidelines and samples with NSC less than 1.1 or RSC less than 1 were removed from the analysis.[4] Also, samples with <10 million effective sequence reads were removed. As a result, 4 samples (RA_CD8_H3K27ac, OA_CD8_H3K27ac, OA_IL17_H3K27ac and RA_CD8_H3K4me3) were excluded and the remaining 86 samples were utilized for further analysis.

SEs were identified with the Rank Ordering of Super-Enhancers (ROSE) algorithm (Key resources information) based on the H3K27ac ChIP sequencing signal with default parameters.

Differentially bound peak analysis for each condition was performed with HOMER software (Key resources information) with fold-enrichment threshold of 2 and Poisson enrichment *P* value threshold of 0.0001.

Motif enrichment analysis was performed with HOMER software. As motifs of some TFs associated with SEs were not included in the HOMER database, we customized motif reference with MotifDb software (Key resources information) for SE associated genes only with *homo sapiens* data.

**Library construction for Hi-C**

Purchased RASFs (n = 7, Articular Engineering) were used to generate an *in situ* Hi-C library as previously described with minor modifications.[5] Briefly, SFs ($1 \times 10^5$ cells/well) were seeded with DMEM (10% FBS, 100 µg/mL L-glutamine, 100 U/mL penicillin, 100 µg/mL streptomycin) into 6-well flat-bottom plates and incubated at 37°C, in 5% $CO_2$. After 12 h, either 10 ng/mL TNF-α or 8-mix was added, and the cells were stimulated for an additional 24 h at 37°C, in 5% $CO_2$. Pooled SFs ($2.8 \times 10^6$ cells) were cross-linked with 1% formaldehyde for 10 min at room temperature. The nuclei were permeabilized, and DNA was digested with 100 units of MboI restriction enzyme

(NEB). The ends of restriction fragments were labeled with biotinylated nucleotides (dATP; Invitrogen), and proximity ligation was performed. After reversal of crosslinks, ligated DNA was purified and sheared to a length of roughly 400 base pairs with Covaris S2 (Covaris), at which point ligation junctions were pulled down with streptavidin beads (Invitrogen). Sequencing libraries were prepared with a Nextera Mate Pair Sample Preparation Kit (Illumina), and sequenced using a HiSeq series (read length of 150 bp, paired-end read).

**Bioinformatics analysis of Hi-C data**

Sequencing reads from 3 samples were mapped to human genome assembly hg19/GRCh37 using BWA-mem which is implemented in JUICER software (ver 1.8.9) (Key resources information). Loops were called using HiCCUPS software (Key resources information) implemented in JUICER with resolution of 5 kbp, 10 kbp and 25 kbp and merged loops were used for downstream analysis.

**SNP typing and imputation**

Genomic DNA from whole blood was isolated using QIAamp DNA Blood Midi Kit (Qiagen). Genotyping was performed using Infinium OmniExpressExome BeadChips (Illumina).

Quality control of the genotyping data was performed using PLINK 1.90 (Key resources information), with a SNP call rate > 0.99, HWE < $1 \times 10^{-6}$ and sample call rate > 0.98. For genome-wide imputation, 595693 post-QC SNPs were pre-phased using SHAPEIT (Key resources information) and imputation was performed using IMPUTE2 (Key resources information) with the 1000 Genomes Phase 3 panel as reference. Post-imputation QC was performed using

SNPTEST (Key resources information). Genotyped and imputed autosomal SNPs or indels with MAF ≥ 0.05 were used for cis-eQTL analysis (6124313 variants in total).

**Cis-eQTL analysis**

For cis-eQTL analysis, genes detected in at least half of the samples under at least 1 condition were included. The count per million (CPM) matrix was normalized between samples using TMM as implemented in edgeR software, normalized across samples using an inverse normal transform and normalized using PEER (Key resources information) with 15 hidden confounders, and the residuals were used for analysis. We used QTLtools (Key resources information) conditional pass for tissue-by-tissue eQTL analysis. In addition, to improve analytical ability, we jointly analyzed the RA and OA samples. We performed a meta-analysis across SFs in various stimulatory conditions and PBMC samples for eQTLs by utilizing Meta-Tissue software (Key resources information), a linear mixed model that allows for heterogeneity in effect sizes across conditions. Cis-eQTL analysis was performed for variants with MAF ≥ 0.05 within a 1 Mb window around each gene.

For each eQTL, we estimated the posterior probability that the effect is shared in each tissue (*m*-value) along with tissue-by-tissue eQTL analysis *P* value.

**GWAS enrichment analysis**

In order to calculate GWAS variant enrichment for epigenomic marks, we prepared 10,000 sets of randomly sampled variants that were matched to GWAS variants for distance from the nearest TSS, minor allele frequency

(MAF), gene density and the number of LD variants ($r^2 \geq 0.5$) using SNPSNAP (Key resources information). We counted the number of GWAS variants or randomly selected variants whose LD ($r^2 \geq 0.8$) variants or itself coincided with epigenomic marks. We calculated the empirical *P* value by comparing the number of GWAS variants that tagged epigenomic marks against the number of randomly selected variants that tagged epigenomic marks. We pruned GWAS variants such that no 2 variants were within 1 Mb of one another, and all GWAS variants within the extended MHC region (25 - 35 Mb on chromosome 6) were removed from the analysis.

**Population enrichment score**

In order to assess the abundance of SF populations reported in the single-cell transcriptome based analysis,[6] we calculated the population enrichment score of each SFs sample. We used "top 20 marker genes for each single-cell RNA sequencing cluster" of 4 SF clusters from the article and calculated the enrichment score of these gene sets using GSVA software (Key resources information) with normalized CPM.

**cDNA synthesis and qRT-PCR**

Purified RASFs ($2 \times 10^4$ cells/well) were seeded with DMEM (10% FBS, 100 µg/mL L-glutamine, 100 U/mL penicillin, 100 µg/mL streptomycin) into 24-well flat-bottom plates and incubated at 37°C, in 5% $CO_2$. After 12 h, 100 U/mL IFN-α, 200 U/mL IFN-γ, 10 ng/mL TNF-α, 10 ng/mL IL-1β, 200 ng/mL IL-6/sIL-6R, 10 ng/mL IL-17, 10 ng/mL TGF-β1, 100 ng/mL IL-18, or 8-mix was added. The cells were stimulated for an additional 3, 10, 24 or 48 h at 37°C, in 5% $CO_2$. In *in vitro* inhibition studies of Brd4 or MTF-1, the cells were pretreated with JQ1 (Sigma; 50 -

5000 ng/mL) or APTO-253 (Medchemexpress; 1 - 4 μg/mL) for 6 h, and incubated with 8-mix for an additional 24 h at 37°C, in 5% $CO_2$.

Total RNAs were extracted with RNeasy Micro Kit (Qiagen) and were reverse-transcribed to cDNA with random primers (Invitrogen), dNTP mixture (Takara), ribonuclease inhibitor (Promega) and SuperScript III (Invitrogen). Quantitative real-time PCR (qRT-PCR) was performed using CFX Connect Real-Time PCR Detection System (Bio-Rad) with QuantiTect SYBR Green PCR Kit (Qiagen). The primer pairs used in this study are shown in online supplementary table 3. Relative expression was calculated based on the abundance of control *GAPDH*.

**Human IL-6 and CCL5 ELISA**

The concentration of IL-6 and CCL5 in supernatants of SFs was measured using the Human IL-6 Uncoated ELISA Kit (Invitrogen) and Human CCL5/RANTES Quantikine ELISA Kit (R & D), respectively, according to the manufacturer's instruction.

**Knockdown assay**

Purchased RASFs were used for knockdown assays. Cells ($4 \times 10^5$ cells/target) were transfected with 300 nM ON-TARGET plus siRNA targeting *MTF1*, *RUNX1*, *SNAI1* or *TCF4* (all from Dharmacon) using a Human Dermal Fibroblast Nucleofector Kit (Lonza) according to the manufacturer's instructions. SiGENOME Non-Targeting Control Pool (300 nM, Dharmacon) was used as a transfection control. Transfected cells ($4 \times 10^4$ cells/well) were seeded with DMEM (10% FBS, 100 μg/mL L-glutamine, 100 U/mL penicillin, 100 μg/mL streptomycin) into 24-

well flat-bottom plates and incubated at 37°C, in 5% $CO_2$. After 6 h, cells were stimulated with 8-mix cytokines for an additional 6 h. Total RNA was isolated using AllPrep DNA/RNA/miRNA Universal Kit. Libraries for RNA sequencing were prepared using TruSeq Stranded mRNA Library Prep Kit. mRNA sequencing was carried out on Illumina MiSeq (read length of 150 bp, paired end).

**CD40 stimulation assay**

Purchased RASFs (n = 3) were used in the CD40 stimulation assay. RASFs ($2 \times 10^4$ cells/well) were seeded with DMEM (10% FBS, 100 µg/mL L-glutamine, 100 U/mL penicillin, 100 µg/mL streptomycin) into 24-well flat-bottom plates and incubated at 37°C, in 5% $CO_2$. After 12 h, cells were stimulated with 1 - 10 ng/mL CD40L (ENZ) and 200 U/mL IFN-γ or 8-mix for an additional 24 h. RNA extraction, cDNA synthesis and RT-PCR was performed as described above. Total RNA was isolated using AllPrep DNA/RNA/miRNA Universal Kit. Libraries for RNA sequencing were prepared using TruSeq Stranded mRNA Library Prep Kit. The mRNA sequencing was carried out on Illumina MiSeq (read length of 150 bp, paired end).

**Mice and Induction of CIA**

DBA/1J male mice (6 weeks) were purchased from Japan SLC. To induce CIA, bovine type II collagen (CII; Chondrex) was emulsified in equal volume of Complete Freund's Adjuvant (CFA; Chondrex). Mice were intradermally injected with the emulsion (100 µL) containing 100 µg CII. Twenty-one days later, a secondary injection was given at the same concentration of CII emulsified in Incomplete Freudn's Adjuvant (IFA; Chondrex).

Mice were examined every day for clinical signs of arthritis after the first injection. The severity of arthritis was assessed by qualitative clinical score determined as follows: 0 = normal paw; 1 =one toe inflamed and swollen; 2 = >1 toe, but not entire paw inflamed and swollen, or mild swelling of entire paw; 3 = entire paw inflamed and swollen; 4 = very inflamed and swollen or ankylosed paw. Each paw was scored individually, and totaled them for each mouse (a maximum of 16 points). All procedures were performed in accordance with National Institutes of Health (NIH) Guide for the Care and Use of Laboratory Animals and approved by the ethics committee of the University of Tokyo Institutional Animal Care and Use Committee.

**Treatment of CIA**

In the treatment experiment, following the onset of arthritis (clinical score >1), CIA mice were randomized into 2 groups (n=7 per each) and intravenously injected with either control (10% DMSO and 18% SBE-β-CD) or 15 mg/kg APTO-253 in 10% DMSO and 18% SBE-β-CD for twice per day for 2 consecutive days per week for 14 days. In the prophylaxis experiment, on the same day as the second immunization, CIA mice were randomized into 2 groups (n=10 per each) and intravenously injected with either control (10% DMSO and 18% SBE-β-CD) or 15 mg/kg APTO-253 in 10% DMSO and 18% SBE-β-CD for twice per day for 2 consecutive days per week for 14 days.

**Histological assessment**

Mice were killed on day 15 after the start of treatment. Four paws were surgically removed and fixed in 4% paraformaldehyde, decalcified in 20% EDTA, and embedded in paraffin. Then, the sectioned tissues were stained with haematoxylin and eosin (H&E) and Safranin O for histopathology. Synovial tissue thickening, mononuclear cell infiltration, pannus invasion, and cartilage damage were graded. For synovial tissue thickening, scores were: 0, no changes (thickness less than 0.7 mm at the patellar tendon); 1; mild changes (0.7-1.0 mm); 2, moderate changes (1.0-2.0 mm); 3, severe changes (2.0 mm and more). For mononuclear cell infiltration, scores were: 0, no changes (no infiltration); 1, mild changes (less than 150 cells in 0.07μm2); 2, moderate changes (150-300 cells); 3, severe changes (300 cells or more). For pannus invasion, scores were: 0, no pannus; 1, mild changes (pannus invasion within the cartilage); 2, moderate changes (pannus invasion into the cartilage/subchondral bone transition); 3, severe changes (pannus invasion into the subchondral bone). For cartilage damage, scores were: 0, no destruction; 1, minimal erosion limited to single spots; 2, slight to moderate erosion in a limited area; 3, more extended erosions; 4, general destruction.

**Statistical Analysis**

For *in vitro* analysis, statistical significance and analysis of variance (ANOVA) between indicated groups were analyzed by R (ver 3.4.1). A comparison of more than 2 group means was analyzed by Tukey's multiple comparison tests. A comparison of 2 group means was analyzed by paired t-test or Mann–Whitney U test. Statistically significant differences were accepted at $P < 0.05$ for all tests. Data in the bar charts were expressed as means ± standard errors of the means (SEM).

**Key resources information**

| REAGENT or RESOURCE | SOURCE | IDENTIFIER |
|---|---|---|
| ***Antibodies*** | | |
| Rabbit polyclonal anti- H3K4me1 | Active Motif | Cat#39298; RRID: AB_2615075 |
| Rabbit polyclonal anti- H3K4me3 | Active Motif | Cat#39160; RRID: AB_2615077 |
| Rabbit polyclonal anti- H3K27ac | Active Motif | Cat#39134; RRID: AB_2561016 |
| Mouse monoclonal anti- CD3 (clone: UCHT1) | BioLegend | Cat#300419; RRID: AB_439780 |
| Mouse monoclonal anti- CD4 (clone: OKT4) | BioLegend | Cat#317433; RRID: AB_11150413 |
| Mouse monoclonal anti- CD8 (clone: RPA-T8) | BioLegend | Cat#301007; RRID: AB_314125 |
| Mouse monoclonal anti- CD14 (clone: M5E2) | BioLegend | Cat#301823; RRID: AB_893253 |
| Mouse monoclonal anti- CD19 (clone: HIB19) | BioLegend | Cat#302211; RRID: AB_314241 |
| Mouse monoclonal anti- CD56 (clone: HCD56) | BioLegend | Cat#318331; RRID: AB_10898118 |
| Mouse monoclonal anti- CD90 (Thy1) (clone: 5E10) | BioLegend | Cat#328107; RRID: AB_893438 |
| ***Biological Samples*** | | |
| Synovial specimen of human rheumatoid arthritis | This paper | N/A |
| Synovial specimen of human osteoarthritis | This paper | N/A |
| Peripheral blood of human rheumatoid arthritis | This paper | N/A |
| Peripheral blood of human osteoarthritis | This paper | N/A |
| ***Chemicals, Peptides, and Recombinant Proteins*** | | |
| Human IFN-α 2A | HumanZyme | Cat#HZ-1066; GenPept: P01563 |
| Human IFN-γ | PeproTech | Cat#300-02; GenPept: P01579 |
| Human TNF-α | PeproTech | Cat#300-01A; GenPept: P01375 |
| Human IL-1β | PeproTech | Cat#200-01B; GenPept: P01584 |

| | | |
|---|---|---|
| Human IL-6 | PeproTech | Cat#200-06; GenPept: P05231 |
| Human sIL-6R | PeproTech | Cat#200-06R; GenPept: P08887 |
| Human TGF-β1 | R&D | Cat# 240-B-002/CF; GenPept: P01137 |
| Human IL-17 | PeproTech | Cat#200-17; GenPept: Q16552 |
| Human IL-18 | MBL | Cat#B001-5; GenPept: Q14116 |
| JQ1 | Sigma-Aldrich | Cat#SML1524; CAS: 1268524-70-4 |
| MEGACD40L | ENZ | Cat#ALX-522-110-C010 |
| APTO-253 | Medchemexpress | Cat#HY-16291; CAS: 916151-99-0 |
| Bovine Type II Collagen | Chondrex | Cat#20022 |
| Complete Freund's Adjuvant | Chondrex | Cat#7001 |
| Incomplete Freund's Adjuvant | Chondrex | Cat#7002 |
| ***Critical Commercial Assays*** | | |
| AllPrep DNA/RNA/miRNA Universal Kit | QIAGEN | Cat#80224 |
| RNeasy Micro Kit | QIAGEN | Cat#74004 |
| TruSeq Stranded mRNA Library Prep Kit | Illumina | Cat#RS-122-2101/2102 |
| ChIP-IT High Sensitivity | Active Motif | Cat#53040 |
| ChIP-IT PBMC | Active Motif | Cat#53042 |
| TruSeq ChIP Library Preparation Kit | Illumina | Cat#IP-202-1012/1024 |
| Nextera Mate Pair Sample Preparation Kit | Illumina | Cat#FC-132-1001 |
| Human Dermal Fibroblast Nucleofector Kit | Lonza | Cat#VPD1001 |
| Human IL-6 Uncoated ELISA Kit | Invitrogen | Cat#88-7066 |
| Human CCL5/RANTES Quantikine ELISA Kit | R&D | Cat#DRN00B |
| ***Deposited Data*** | | |
| Read counts data of RNA sequencing | This paper | NBDC: hum0207.v1.RNA.v1 |

| | | |
|---|---|---|
| eQTL summary | This paper | NBDC: hum0207.v1.eQTL.v1 |
| Peaks data of ChIP sequencing | This paper | NBDC: hum0207.v1.ChIP.v1 |
| Chromatin loops data of Hi-C | This paper | NBDC: hum0207.v1.HiC.v1 |
| ***Experimental Models: Cell Lines*** | | |
| Synovial fibroblasts of human rheumatoid arthritis | Articular Engineering | Cat#CDD-H-2910-RA |
| ***Experimental Models: Organisms/Strains*** | | |
| Mouse: DBA/1J Jms Slc | SLC | RRID: MGI:5651994 |
| ***Oligonucleotides*** | | |
| ON-TARGET plus siRNA targeting MTF1 | Dharmacon | Cat#L-020078-00-0005 |
| ON-TARGET plus siRNA targeting RUNX1 | Dharmacon | Cat#L-003926-00-0005 |
| ON-TARGET plus siRNA targeting SNAI1 | Dharmacon | Cat#L-010847-01-0005 |
| ON-TARGET plus siRNA targeting TCF4 | Dharmacon | Cat#L-004594-00-0005 |
| SiGENOME Non-Targeting Control Pool #1 | Dharmacon | Cat#D-001206-13-05 |
| ***Software and Algorithms*** | | |
| R3.4.1 | R Core team, 2016 | https://www.R-project.org |
| Python 2.7 | Python Software | https://www.python.org/ |
| STAR (version 2.5.3) | Dobin et al, 2012 | https://github.com/alexdobin/STAR |
| HTSeq (version 0.6.0) | Anders et al, 2014 | https://github.com/simon-anders/htseq |
| RSEM (version 1.3.0) | Li et al, 2011 | https://github.com/deweylab/RSEM |

| | | |
|---|---|---|
| RUVseq (version 1.10.0) | Risso et al, 2014 | https://github.com/drisso/RUVSeq |
| edgeR (version 3.18.1) | Robinson et al, 2010 | https://bioconductor.org/packages/release/bioc/html/edgeR.html |
| MAGMA (version 1.07) | de Leeuw C et al, 2015 | https://ctg.cncr.nl/software/magma |
| Bowtie2 (version 2.3.4.2) | Langmead et al, 2009 | https://github.com/BenLangmead/bowtie2 |
| MACS 2.0 (version 2.1.1) | Zhang et al, 2008 | https://github.com/taoliu/MACS |
| ROSE | Whyte et al, 2013 | http://younglab.wi.mit.edu/super_enhancer_code.html |
| HOMER (version 4.9.1) | Heinz et al, 2010 | http://homer.ucsd.edu/homer/ |
| MotifDb (version 1.18.0) | Shannon et al, 2019 | http://bioconductor.org/packages/release/bioc/html/MotifDb.html |
| BWA (version 0.7.17) | Li et al, 2009 | https://github.com/lh3/bwa |
| JUICER (version 1.5) | Durand et al, 2016 | https://github.com/aidenlab/juicer |
| HiCCUPS | Rao et al, 2014 | https://github.com/aidenlab/juicer/wiki/HiCCUPS |
| PLINK (v1.90b4.4) | Purcell et al, 2007 | https://github.com/chrchang/plink-ng/ |
| SHAPEIT (v2.r904) | Delaneau et al, 2012 | https://mathgen.stats.ox.ac.uk/genetics_software/shapeit/shapeit.html |
| IMPUTE2 (version 2.3.2) | Howie et al, 2009 | https://mathgen.stats.ox.ac.uk/impute/impute_v2.html |

| | | |
|---|---|---|
| SNPTEST (version 2.5.4) | Marchini et al, 2007 | https://mathgen.stats.ox.ac.uk/genetics_software/snptest/snptest.html |
| Meta-Tissue (version 0.5) | Sul et al, 2013 | http://genetics.cs.ucla.edu/metatissue/ |
| PEER (version 1.0) | Stegle et al, 2010 | https://github.com/PMBio/peer/wiki |
| QTLtools | Delaneau, 2017 | https://qtltools.github.io/qtltools/ |
| SNPSNAP | Pers et al, 2014 | https://data.broadinstitute.org/mpg/snpsnap/ |
| GSVA (version 1.24.2) | Hänzelmann et al, 2013 | https://bioconductor.org/packages/release/bioc/html/GSVA.html |
| BEDTools (version 2.26.0) | Quinlan et al, 2010 | https://github.com/arq5x/bedtools2 |
| ***Other*** | | |
| Human CD14 MicroBeads | Miltenyi Biotec | Cat#130-050-201 |
| Mbo Ⅰ | NEB | Cat#R0147S |
| Biotin-14-dATP | Invitrogen | Cat#19524016 |
| DynabeadsTM MyOne™ Streptavidin T1 | Invitrogen | Cat#65601 |
| SuperScript III Reverse Transcriptase | Invitrogen | Cat#18080093 |
| Random Primers | Invitrogen | Cat#48190011 |
| dNTP Mixture | Takara | Cat#4030 |
| RNasin® Plus Ribonuclease Inhibitor | Promega | Cat#N2611 |
| QuantiTect SYBR Green PCR Kit | QIAGEN | Cat#204143 |

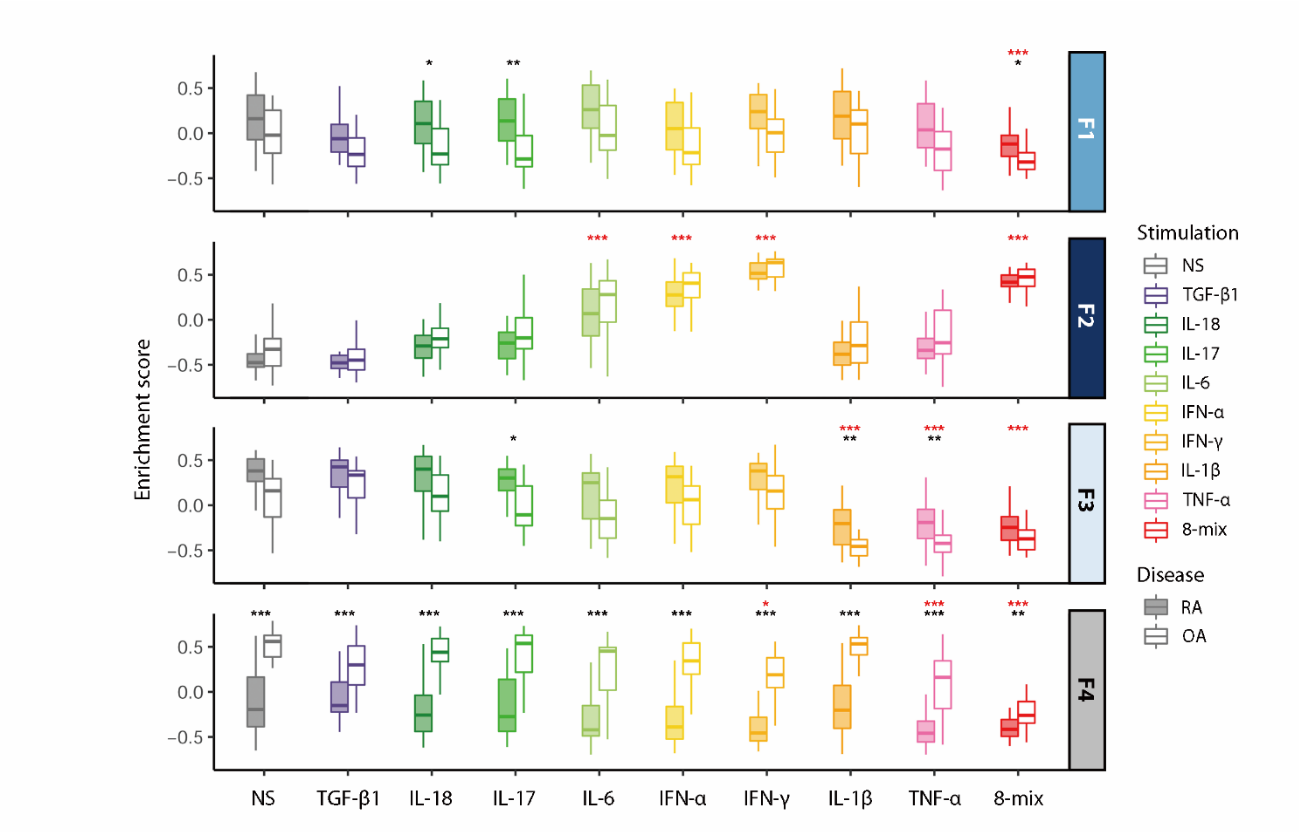


**Online supplementary figure 1. The estimated enrichment of SF populations revealed by single-cell RNA sequencing analysis of freshly isolated SFs.**

The population enrichment score of our stimulated SFs was calculated using GSVA software. Fan Zhang *et al* reported three sublining subsets, $CD34^{+}$ (F1), $HLA\text{-}DRA^{hi}$ (F2), and $DKK3^{+}$ (F3) and one lining subset (F4) based on single-cell RNA sequencing.[6]

We applied "top 20 marker genes for each single-cell RNA sequencing cluster" for enrichment score calculation. $P$ values were calculated using a paired t test (*$P < 0.05$, **$P < 0.001$, ***$P < 0.0001$, red, comparison between stimulation and non-stimulation; black, comparison between RA and OA). Boxes, interquartile range; whiskers, distribution.

RA, rheumatoid arthritis; OA, osteoarthritis; NS, non-stimulation; F1-4, Fraction 1-4.

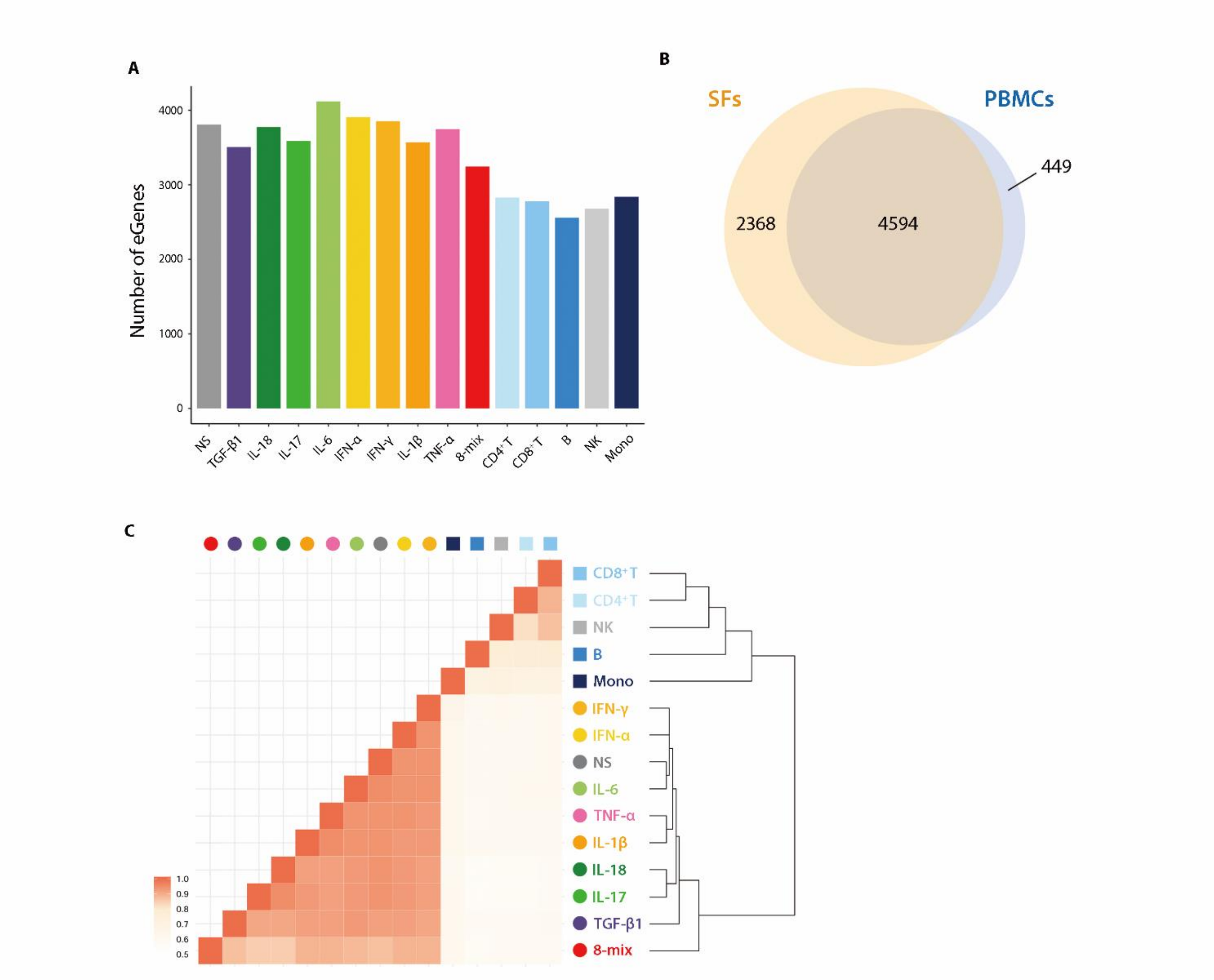


**Online supplementary figure 2. Summary of the result of cis-eQTL analysis in stimulated SFs and PBMCs from RA and OA patients.**

(**A**) The number of eGenes in stimulated SFs and PBMCs. (**B**) A Venn diagram representing the overlap of eGenes in SFs (in any condition) and PBMCs. (**C**) The correlation of cis-eQTL meta-analysis effect size between each stimulated SFs and PBMCs.

Spearman's ρ is shown in heatmap. Samples were hierarchically clustered according to correlation.

NS, non-stimulation; SF, synovial fibroblast; PBMC, peripheral blood mononuclear cell; eQTL, expression quantitative trait locus.

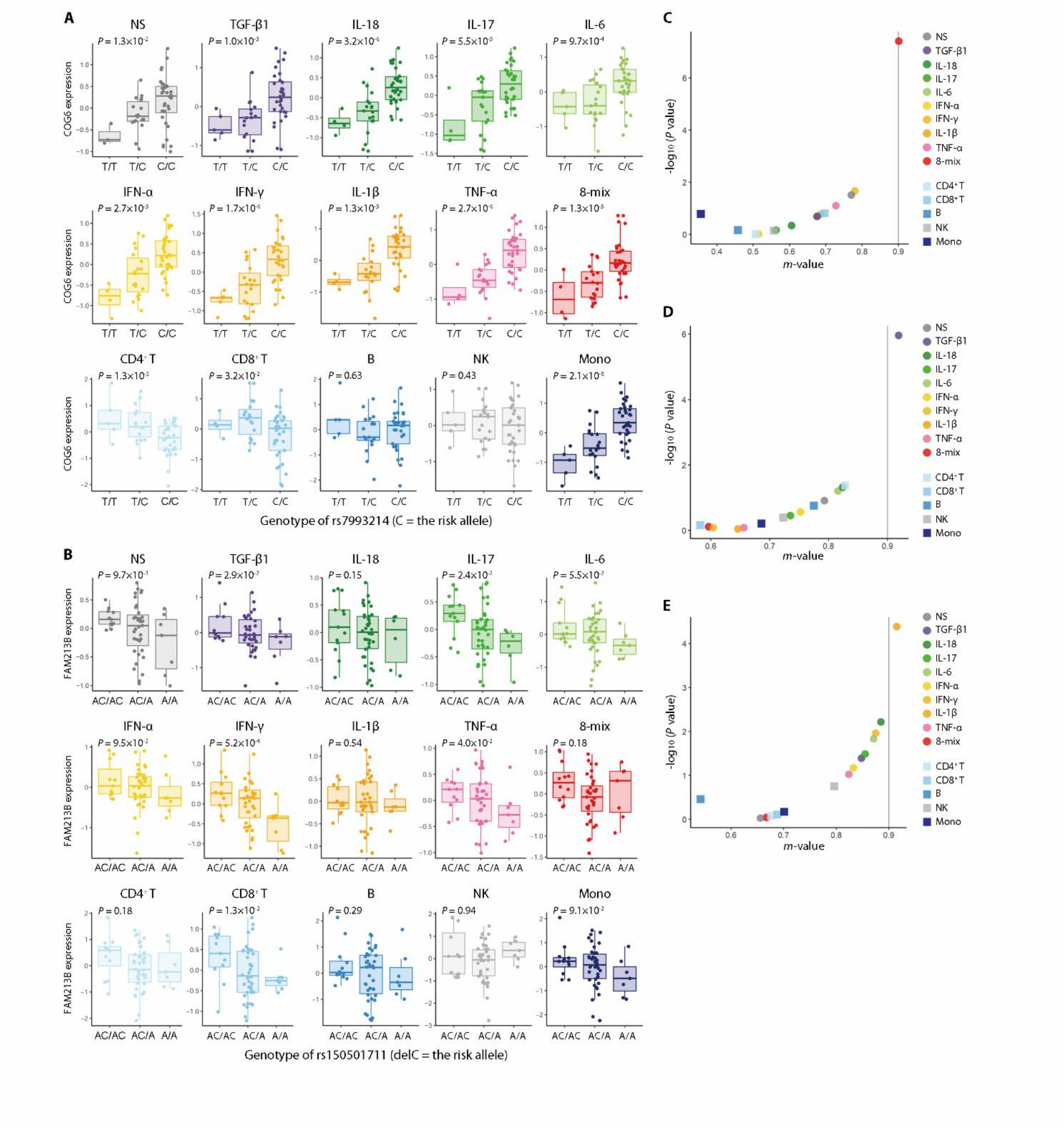


**Online supplementary figure 3. Representative examples of eQTL in stimulated SFs.**

(**A, B**) Boxplot of eQTLs which showed possible colocalization with RA GWAS top-associated loci (also see online supplementary table 1). *COG6* normalized expression stratified by rs7993214 genotype (**A**) and *FAM213B* normalized expression stratified by rs150501711 genotype (**B**). Nominal *P* values of eQTL analysis are shown. (**C** to **E**) Examples of stimulation-specific eQTLs. A dot plot of cis-eQTL meta-analysis posterior probability M value and tissue-by-tissue analysis -$\log_{10}$ *P* value for rs8069701-*CD300E* (**C**), chr1:160179300-*SLAMF1* (**D**) and rs914744-*TRAF2* (**E**) are shown.

Boxes, interquartile range; whiskers, distribution; dots, outliers. NS, non-stimulation; eQTL, expression quantitative trait locus.

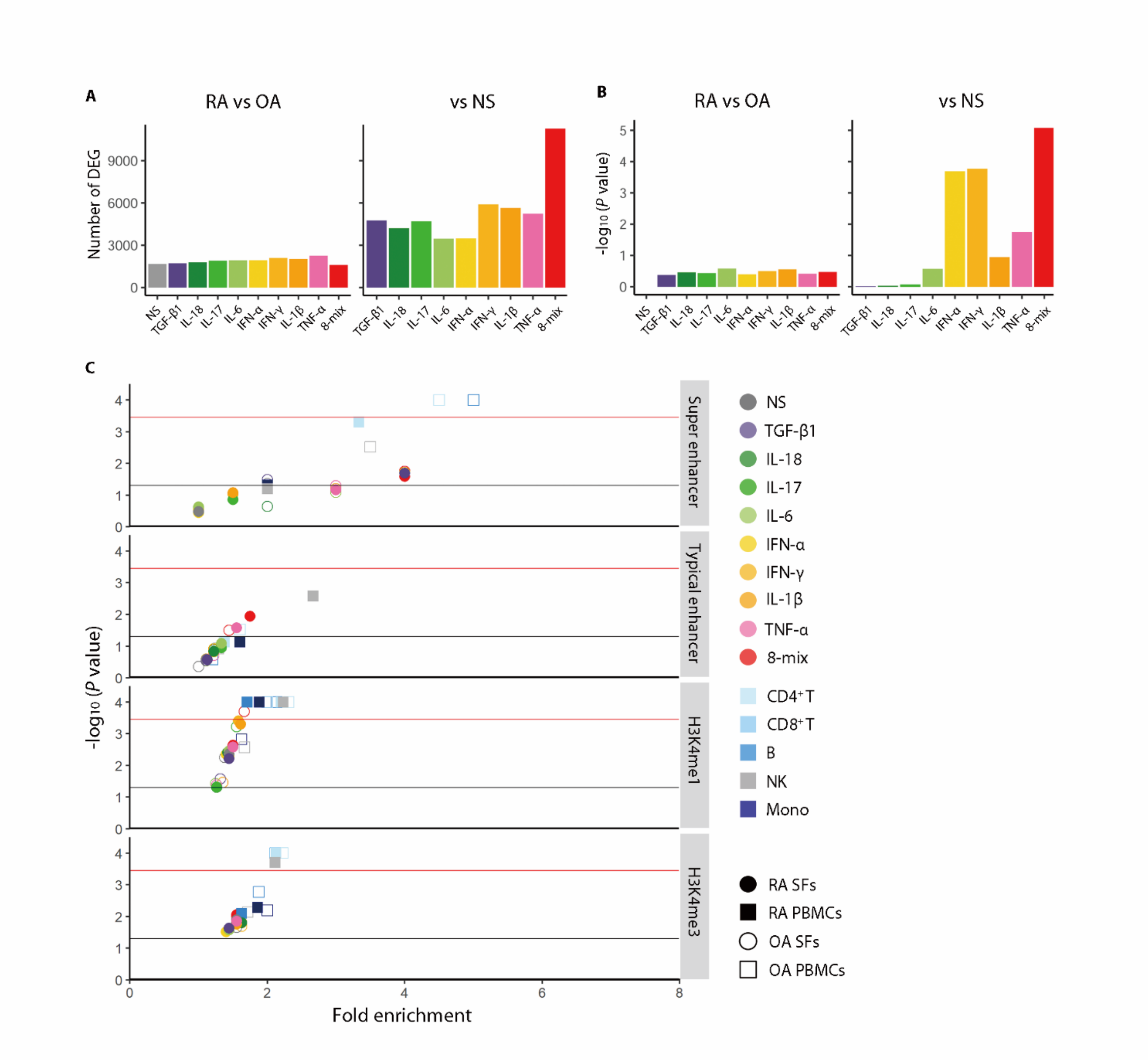


**Online supplementary figure 4. Polygenic association between differentially expressed genes and RA risk loci, and GWAS-super-enhancers enrichment analysis in the other immune related disease.**

(**A**) The number of differentially expressed genes from RNA sequencing. Left, RA vs OA; Right, stimulated SFs vs unstimulated SFs. (**B**) Polygenic association analysis of differentially expressed genes with RA genetic risk. Association *P* values were calculated with MAGMA. (**C**) Enrichment of type 1 diabetes mellitus risk loci in transcriptional regulatory regions of stimulated SFs and PBMCs. Active enhancers were classified into super-enhancers (SEs) and typical-enhancers (TEs) following standard ROSE algorithms. The red solid lines ($-\log_{10}$ (*P* value) = 3.2) and the black solid lines ($-\log_{10}$ (*P* value) = 1.3) are the cutoff for Bonferroni significance and nominal *P* = 0.05, respectively.

DEG, differentially expressed genes; RA, rheumatoid arthritis; OA, osteoarthritis; NS, non-stimulation; SE, super enhancer; TE, typical enhancer.

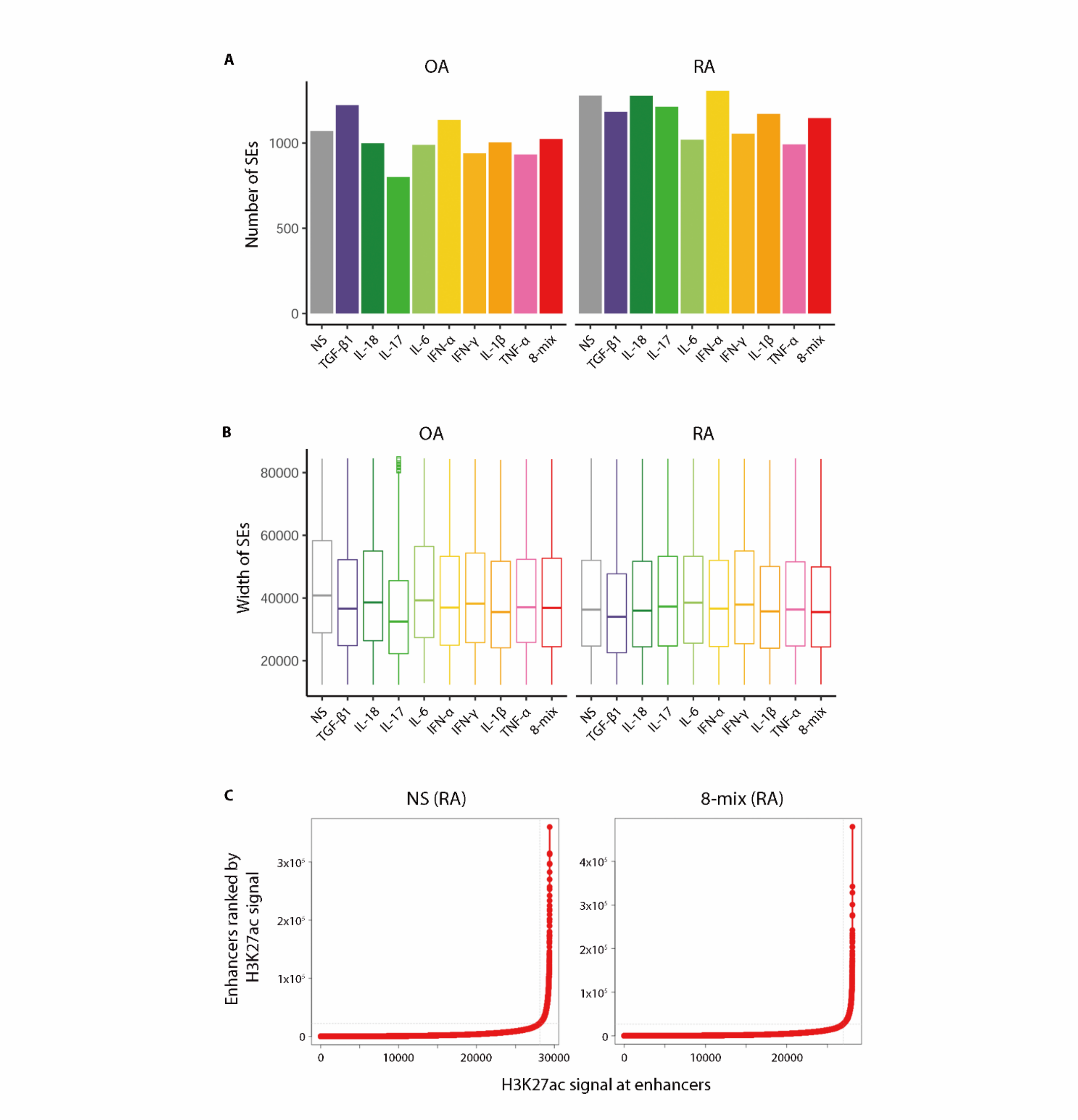


**Online supplementary figure 5. Comparison of the number and width of SEs between stimulatory conditions of SFs.**

(**A**) The number of SEs of each stimulated SFs. (**B**) The width of SEs of each stimulated SFs. Boxes, interquartile range; whiskers, distribution. (**C**) Distribution of SEs based on H3K27Ac signals and enhancer ranks in nonstimulating (left) and 8-mix stimulating (rignt) RASFs.

RA, rheumatoid arthritis; OA, osteoarthritis; NS, non-stimulation; SE, super enhancer.

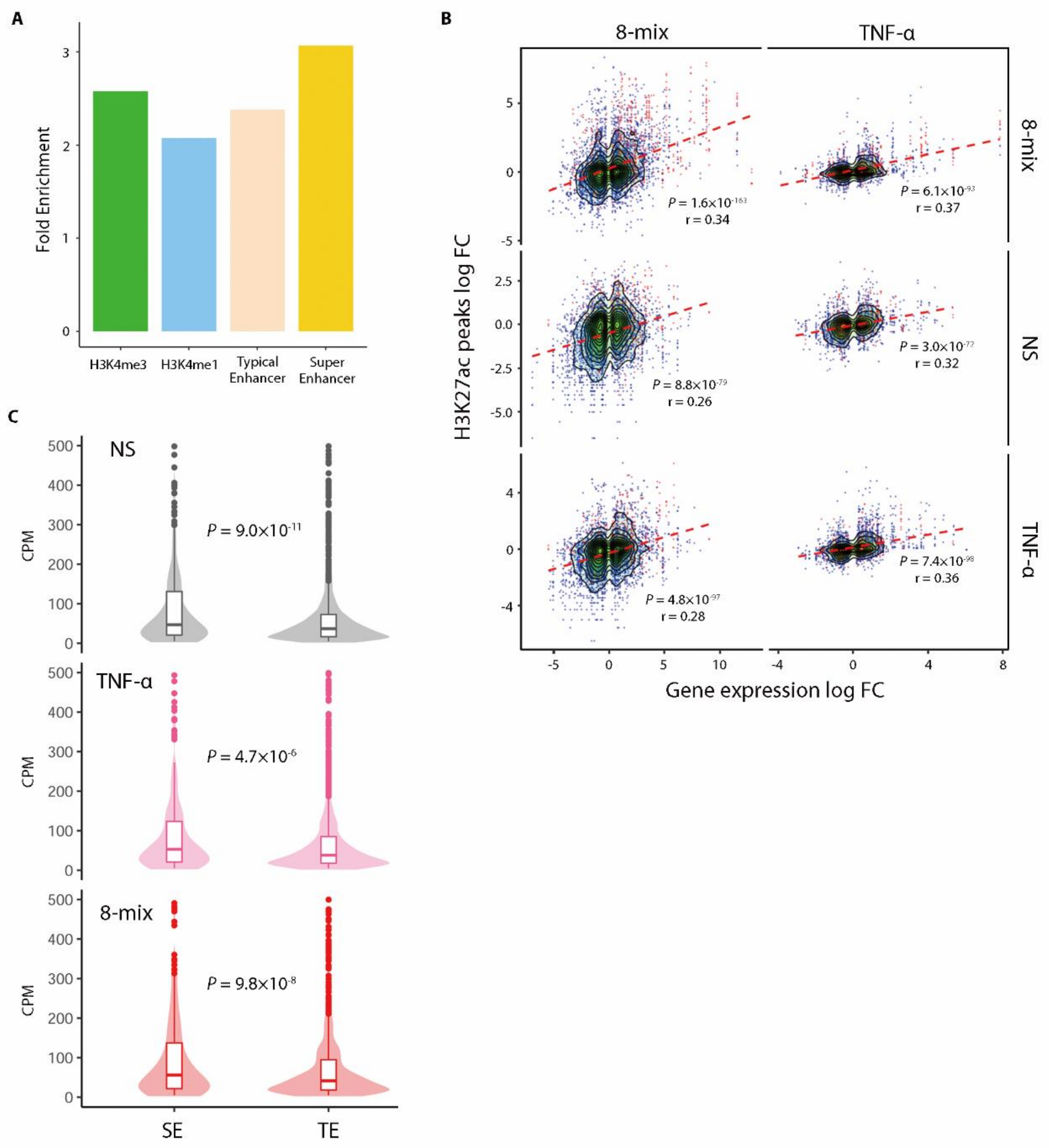


**Online supplementary figure 6. The relevance among Hi-C loop, SEs and gene expression levels.**

(**A**) The enrichment of Hi-C loop anchors with epigenome status (H3K4me3, H3K4me1, TEs and SEs). Fold change was calculated compared with randomly selected loci from the whole genome. (**B**) Comparison of H3K27ac peak fluctuation and its associated gene expression fluctuation. Associated gene for each H3K27ac peak was determined with Hi-C loops. Fold change between RA and OA samples are compared. In the left panel, genes and

H3K27ac peaks were connected with Hi-C loops in 8-mix stimulated SFs. In the right panel, genes and H3K27ac peaks were connected with Hi-C loops in TNF-α stimulated SFs. Top, comparison in 8-mix stimulating SF samples; middle, comparison in unstimulated SF samples; bottom, comparison in TNF-α stimulating SF samples. *P* values and ρ were calculated using Spearman's test. Red dot, SE-contacting genes; Blue dot, TE-contacting genes. (**C**) Transcript abundances of TE or SE-contacting genes from RNA sequencing data for stimulated SFs. Boxes, interquartile range; whiskers, distribution; dots, outliers; density plot, frequency. *P* values were calculated with Student t test.

NS, non-stimulation; SE, super enhancer; TE, typical enhancer; CPM, count per million.

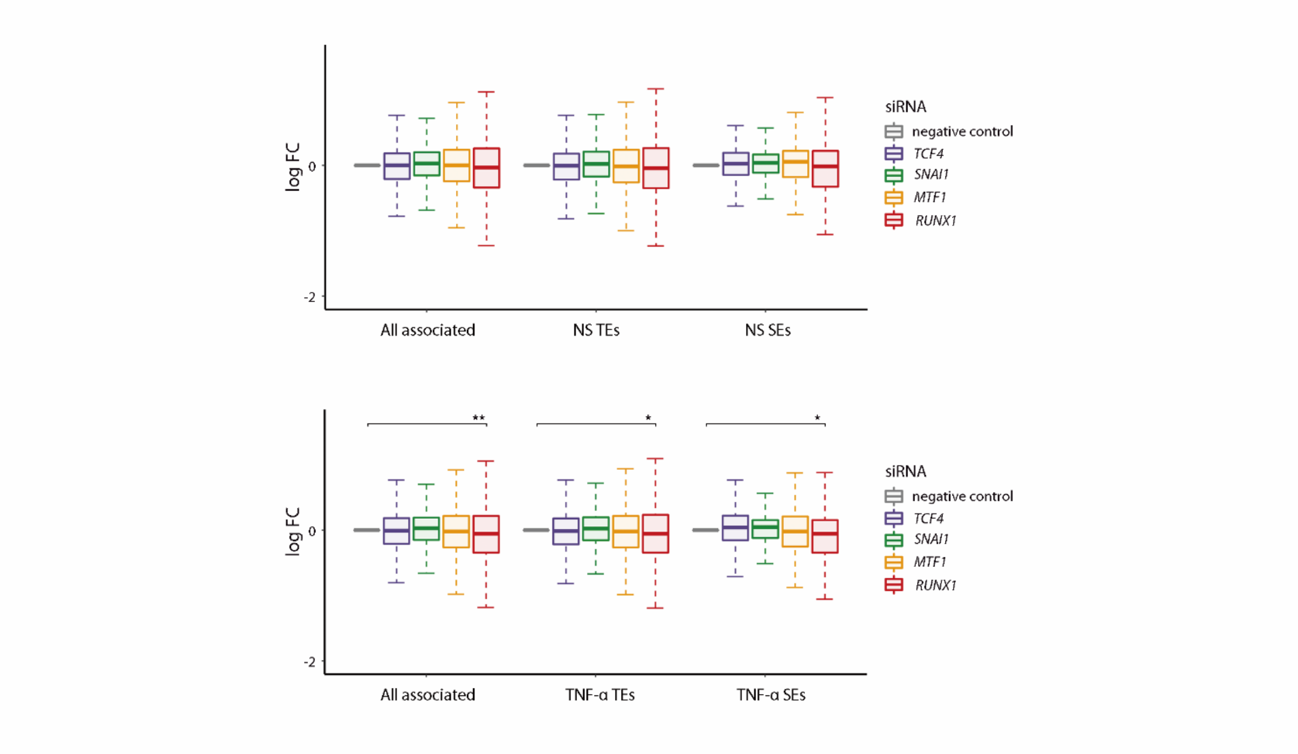


**Online supplementary figure 7. Transcription factors associated with SEs formation in 8-mix SFs.**

Expression of non-stimulation or TNF-α TE or SE-contacting genes in transcription factors (*TCF4*, *SNAI1*, *MTF1* and *RUNX1*) -depleted SFs relative to control SFs. Boxes, interquartile range; whiskers, distribution. *P* values were calculated using a paired t test (**P* < 0.05, ***P* < 0.001).

NS, non-stimulation; SE, super enhancer; TE, typical enhancer.

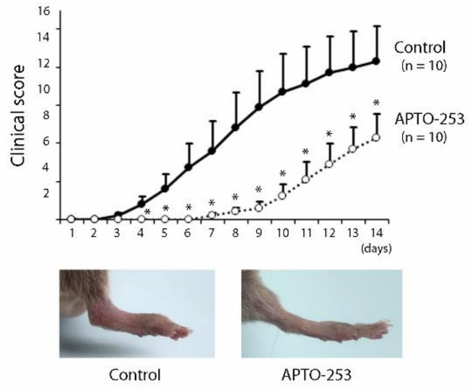


**Online supplementary figure 8. Preventive effect of APTO-253 on CIA model.**

On the same day as the second immunization, CIA mice were intravenously injected with either control or 15 mg/kg APTO-253 for twice per day for 2 consecutive days per week. Clinical scores in each group and representative pictures of hind paw. Dots, mean; Error bars, SEM. *P* values were calculated using a Mann–Whitney U test (*, $P < 0.05$).

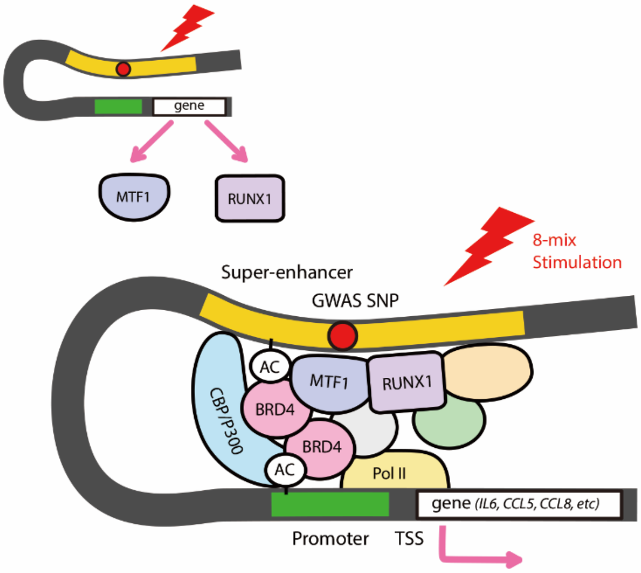


**Online supplementary figure 9. A Graphical summary of the present study.**

The authors conducted integrated analysis of activated SFs, and demonstrated chromatin remodeling in the presence of synergistic proinflammatory cytokines. The dynamic changes with super-enhancer formation were associated with RA heritability. Some transcription factors (MTF1, RUNX1) could be crucial for this structural rearrangement and the formation of inflammatory arthritis.

TSS, transcriptional start site.

**Online supplementary table 1. Colocalization of RA GWAS and cis-eQTL signals (Separate file).**

The list of cis-eQTL top variants which are in LD with RA GWAS top-associated loci. Variant pairs with $R^2 > 0.6$ in EUR or EAS population in 1000G phase 3 data are listed. The RA GWAS top-associated SNP was downloaded from the NHGRI-EBI GWAS Catalog.[7] EFO_0000685 was downloaded on 11/09/2018.

eQTL, expression quantitative trait locus; GWAS, lead variant in genome-wide association study; $R^2$, r square values between eQTL top variant and GWAS top-associated SNP in EUR or EAS population (the larger one is written); method, eQTL analysis method for the indicated top eQTL variant; MT, meta-tissue analysis; TBT, tissue-by-tissue analysis

**Online supplementary table 2. Summary of SE-contacted genes (Separate file).**

**Online supplementary table 3. Sequences of primer pairs used in qRT-PCR.**

| Target | Sequence (Forward) | Sequence (Reverse) |
|---|---|---|
| *IL6* | 5'- GCCTTCGGTCCAGTTGCCTT -3' | 5'- AGTGCCTCTTTGCTGCTTTCAC -3' |
| *CCL5* | 5'- AGTGTGTGCCAACCCAGAGAAGAA -3' | 5'- TGTGGTAGAATCTGGGCCCTTCAA -3' |
| *MTF1* | 5'- GCCCCAGTAATGGCTGTGAG -3' | 5'- TCCTCTGATCCATTGTGTTGTGG -3' |
| *RUNX1* | 5'- TCCACTGCCTTTAACCCTCA -3' | 5'- AGGTGAAATGGGCGTTGCT -3' |
| *SNAI1* | 5'- TATGCTGCCTTCCCAGGCTTG -3' | 5'- ATGTGCATCTTGAGGGCACCC -3' |
| *TCF4* | 5'- TCCAGGTTTGCCATCTTCAGT -3' | 5'- GCCTGGCGAGTCCCTATTG -3' |
| *GAPDH* | 5'- GAAGGTGAAGGTCGGAGTC -3' | 5'- GAAG ATGGTGATGGGATTTC -3' |

**Online supplementary table 4. Characteristics of patients for RNA sequencing.**

| | RA (n=30) | OA (n=30) |
|---|---|---|
| **Age, years (range)** | 70 (52-80) | 73 (54-88) |
| **Female, n (%)** | 26 (86.7) | 24 (80.0) |
| **Serological markers** | | |
| Rheumatoid factor-positive, n (%) | 25 (83.3) | 0 (0) |
| Anti-CCP autoantibody-positive, n (%) | 21 (70.0) | 0 (0) |
| **Disease activity, n (%)** | | |
| High (DAS-ESR >5.1) | 6 (20.0) | n/a |
| Moderate (DAS-ESR 3.2-5.1) | 20 (66.7) | n/a |
| Low/Remission (DS-ESR <3.2) | 4 (13.3) | n/a |
| **Steinbrocker Stage, n (%)** | | |
| Ⅰ | 0 (0) | n/a |
| Ⅱ | 3 (10.0) | n/a |
| Ⅲ | 11 (36.7) | n/a |
| Ⅳ | 16 (53.3) | n/a |
| **Steinbrocker Class, n (%)** | | |
| Ⅰ | 0 (0) | n/a |
| Ⅱ | 17 (56.7) | n/a |
| Ⅲ | 11 (36.7) | n/a |
| Ⅳ | 2 (6.7) | n/a |
| **Treatment, n (%)** | | |
| csDMARDs | | |
| MTX | 14 (46.7) | 0 (0) |
| BUC | 5 (16.7) | 0 (0) |
| IGU | 4 (13.3) | 0 (0) |
| SASP | 5 (16.7) | 0 (0) |
| TAC | 5 (16.7) | 0 (0) |
| Glucocorticoids | 21 (70.0) | 0 (0) |
| boDMARDs | | |
| TNFi | 6 (20.0) | 0 (0) |
| ABT | 2 (6.7) | 0 (0) |
| TCZ | 2 (6.7) | 0 (0) |

RA, rheumatoid arthritis; OA, osteoarthritis; csDMARDs, conventional synthetic disease-modifying antirheumatic drugs; MTX, methotrexate; BUC, bucillamine; IGU, iguratimod; SASP, salazosulfapyridine; TAC, tacrolimus;

boDMARDs, biological originator disease-modifying antirheumatic drugs; TNFi, tumour necrosis factor inhibitors;

ABT, abatacept; TCZ, tocilizumab; n/a, not applicable.
